# IL-17 undermines longevity and stress tolerance by inhibiting a protective transcriptional network

**DOI:** 10.1101/2023.01.13.523898

**Authors:** Qiongxuan Lu, Ioana Vladareanu, Lina Zhao, Lars Nilsson, Johan Henriksson, Changchun Chen

## Abstract

Aberrant cytokine secretion contributes to the pathogenesis of autoimmune diseases and age-related disorders, but the molecular mechanism underlying this is not entirely clear. Here, we elucidate how interleukin-17 (IL-17) overactivation shortens lifespan and damages defense mechanisms against stress in *C. elegans*. Our analysis reveals that NHR-49, the *C. elegans* ortholog of human PPARα and HNF4, is the central component in the transcriptional network undermined by increased IL-17 signaling. Both NHR-49 and its coactivator MDT-15 physically interact with the downstream components of IL-17 pathway, and their expression is significantly decreased when IL-17 signaling is enhanced. IL-17 overactivation also induces the expression and nucleus entry of the *C. elegans* ortholog of NF-κB inhibitor NFKI-1/IκBζ to repress the activity of transcriptional coactivator MDT-15 and CBP-1. IL-17 signaling acts on neurons to modulate the activity of NFKI-1/IκBζ and NHR-49. In addition, persistent IL-17 activation decreases the expression of HLH-30/TFEB, leading to the reduced transcription of lysosomal lipase genes in the distal tissues. All these jointly contribute to the increased sensitivity to oxidative stress of animals with enhanced IL-17 signaling. Collectively, our work illustrates a transcription system undermined by IL-17 overactivation in the animals without NF-κB, and provides mechanistic insight into the pathogenesis of abnormal IL-17 secretion.

## Introduction

The ability to cope with stress and harmful stimuli is critical to maintain cellular and organismal homeostasis. On the molecular level, cells coordinate protective transcriptional responses and the secretion of immune effectors to remove detrimental threats. Cytokines, a group of small secreted proteins, are the major signaling molecules in defensive response to infections (Kany, Vollrath, & Relja, 2019; Takeuchi & Akira, 2010). Although the release of cytokines is necessary to repair the damages, their aberrant activation often underlies the pathogenesis of autoimmune disorders (Kany et al., 2019; Takeuchi & Akira, 2010). Similarly, the role of cytokines in the nervous system is also bilateral. Basal secretion of proinflammatory cytokines such as interleukin (IL-1) contributes actively to neuronal communications (Nemeth & Quan, 2021). However, pathological upregulation of cytokines directly activates nociceptive neurons, leading to the development of hyperalgesia and the progression of several age-related neurological disorders (Cook, Christensen, Tewari, McMahon, & Hamilton, 2018; Kinney et al., 2018; McCauley & Baloh, 2019).

The interleukin-17 (IL-17) cytokines are critical mediators in innate and adaptive immunity, as well as in the progression of inflammatory diseases such as rheumatoid arthritis (RA) and multiple sclerosis (Gaffen, 2009; Lubberts, 2008; Pappu, Ramirez-Carrozzi, Ota, Ouyang, & Hu, 2010). Accumulating evidence suggest that IL-17 sensitizes nociceptive sensory neurons in the generation of inflammatory pain. For example, the injection of IL-17 effectively induces mechanical hyperalgesia (Ebbinghaus et al., 2017; Luo et al., 2019; Richter et al., 2012), whereas IL-17 neutralization by antibodies significantly attenuates it (Richter et al., 2012). However, relatively less is known about the long-term consequences of persistent IL-17 activation.

In *C. elegans*, IL-17 signaling is required for sustaining the responsiveness of oxygen (O_2_) sensing circuit to the aversive stimulation of 21% O_2_ (Chen et al., 2017; Flynn et al., 2020). Signal transduction is initiated by the binding of ILC-17.1, a homolog of human IL-17s, to the heteromeric receptor complex ILCR-1/ILCR-2 (Chen et al., 2017) (Figure 1A). The receptor complex further recruits a signalosome, which includes the receptor adaptor ACTL-1, PIK-1/IRAK, the scaffold protein MALT-1 and the *C. elegans* ortholog of human NF-κB inhibitor NFKI-1 (Brena et al., 2020; Chen et al., 2017; Flynn et al., 2020) (Figure 1A). Even though the NF-κB inhibitor ortholog exists, *C. elegans* genome does not seem to encode apparent NF-κB-like transcription factors (Irazoqui, Urbach, & Ausubel, 2010). Therefore, how ILC-17.1 signaling modulates transcription remains to be elucidated. Here, we show that increased ILC-17.1 signaling shortens animals’ longevity and impairs their tolerance to oxidative stress, by inhibiting multiple functionally-related transcription factors and regulators. Our observations elucidate a coordinated regulatory network, which shapes the transcriptional outputs of ILC-17.1 signaling in *C. elegans*.

**Figure 1.**
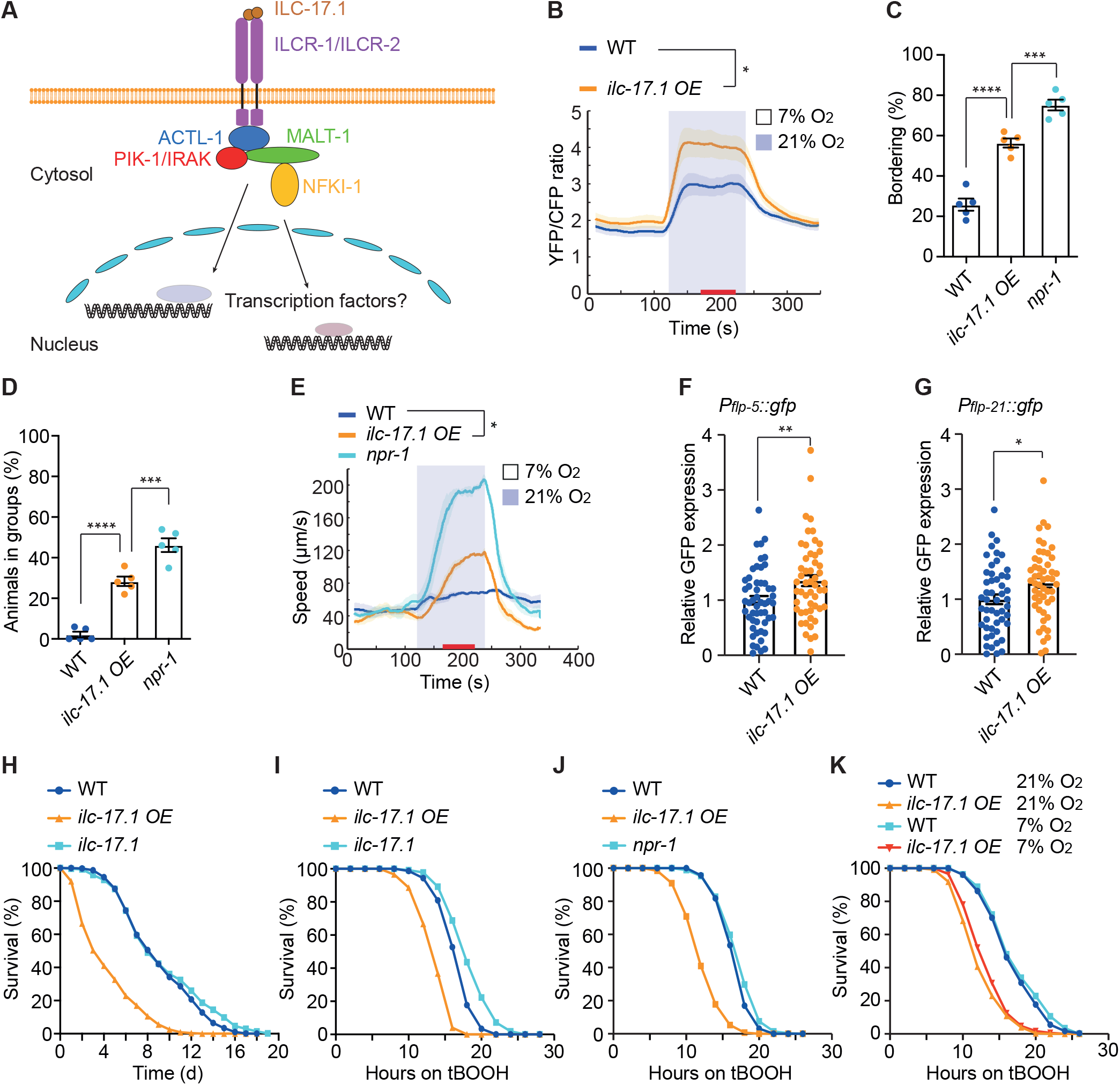
ILC-17.1 overactivation enhances hyperoxia responses and impairs longevity and stress tolerance. **(A)** Schematic of the known components in ILC-17.1 (IL-17) signaling in *C. elegans*. **(B)** O_2_-evoked Ca^2+^ transients in RMG interneurons of WT and *ilc-17*.*1 OE* animals. Red bar on the X axis corresponds to the time window for statistical analysis. *, *p*<0.05, Mann-Whitney U test. (**C** and **D**) Bordering (C) and aggregation (D) assays of indicated genotypes: WT (N2), *ilc-17*.*1* overexpression (OE) and *npr-1(ad609). ilc-17*.*1 OE* was generated by integrating *ilc-17*.*1* extrachromosomal arrays into the genome. Each strain was assayed 5 times with a total number of 300 animals. Error bars indicate standard error of the mean (SEM). ****, *p*<0.0001 and ***, *p*<0.001. ANOVA with Tukey correction. (**E**) Locomotory responses to switches between 7% and 21% O_2_ of indicated genotypes: WT, *ilc-17*.*1 (OE)* and *npr-1(ad609)*. Speed was recorded for 2 mins at each O_2_ interval. Red bar on the X axis corresponds to the time window for statistical analysis. Each strain was assayed at least 3 times with >75 animals. *, *p*<0.05. Mann-Whitney U test. (**F** and **G**) Expression of *Pflp-5::GFP* (F) and *Pflp-21::GFP* (G) in the RMG interneurons of WT (n=52 in F, and n=50 in G) and *ilc-17*.*1 OE* animals (n=52 in F, and n=51 in G). Error bars indicate SEM. **, *p*=0.006; *, *p*=0.018, *t* test. (**H**) Longevity survival curves of WT, *ilc-17*.*1 OE*, and *ilc-17*.*1(tm5218)* animals. In this and all subsequent lifespan assays, day-one adult animals were picked to each assay plate as day 0. At least two independent experiments were included, and each experiment has at least 3 technical replicates. Numerical values and statistical analyses are in Supplementary File 1. (**I**) Survival curves of WT, *ilc-17*.*1 OE*, and *ilc-17*.*1(tm5218)* animals on 7.7mM tBOOH. In this and all subsequent survival assays, synchronized day-one adult animals were picked to each assay plate. At least two independent experiments were included, and each experiment contains at least 3 technical replicates. Numerical values and statistical analyses are in Supplementary File 1. (**J**) Survival curves of WT, *ilc-17*.*1 OE* and *npr-1(ad609)* mutant animals on 7.7mM tBOOH. Numerical values and statistical analyses are in Supplementary File 1. (**K**) Survival curves of WT and *ilc-17*.*1 OE* animals grown and assayed at 21% O_2_ on 7.7mM tBOOH, and of WT and *ilc-17*.*1 OE* animals grown at 7% O_2_ for two generations and assayed at 7% O_2_ on 7.7mM tBOOH. Numerical values and statistical analyses are in Supplementary File 1.

## Results

### Constant activation of ILC-17.1 signaling heightens hyperoxia response and undermines stress tolerance

In mammals, an initial burst of IL-17 is critical for the immune defense against infections (Gaffen, 2009; Lubberts, 2008; Pappu et al., 2010). We wondered if the expression of *ilc-17*.*1* in *C. elegans* could also be induced by infections. It has been reported that *ilc-17*.*1* levels were increased upon exposure to *Enterococcus faecalis* (Engelmann et al., 2011). By using a transcriptional mCherry reporter under the control of *ilc-17*.*1* promoter, we confirmed that the expression of *ilc-17*.*1* was significantly increased after 8 hours of *E. faecalis* exposure, and its expression was continuously induced as long as *E. faecalis* was present (Figure 1–figure supplement 1A and B). Since IL-17 over-secretion contributes to the pain by sensitizing nociceptors (Ebbinghaus et al., 2017; Luo et al., 2019; Richter et al., 2012), we explored if persistent ILC-17.1 expression in *C. elegans* could also modify the nociceptive property. We overexpressed *ilc-17*.*1* (*ilc-17*.*1 OE*) and examined the O_2_ sensing circuit in *C. elegans*. The standard *C. elegans* laboratory strain, N2, fails to aggregate or move rapidly at 21% O_2_ due to a gain of function (*gof*) mutation in *npr-1*. NPR-1 acts downstream of Ca^2+^ transients to reduce the signal output from RMG interneurons (Laurent et al., 2015; Macosko et al., 2009), whereas ILC-17.1 signaling is required for O_2_-evoked Ca^2+^ transients in RMG interneurons (Chen et al., 2017). Consistent with previous observations (Chen et al., 2017), *ilc-17*.*1 OE* heightened the Ca^2+^ transients in RMG interneurons when O_2_ levels were increased from favored 7% to aversive 21% (Figure 1B), and partially restored O_2_-related behaviors to N2 animals (Figure 1C–E), suggesting that ILC-17.1 signaling in *C. elegans* can also modify the excitability of nociceptive circuits.

RMG interneurons release neuropeptides to induce rapid movement at 21% O_2_ (Laurent et al., 2015). RNA sequencing (RNA-seq) analysis showed that the expression of many FMRFamide-like neuropeptides was slightly increased by *ilc-17*.*1 OE*, including *flp-5* and *flp-21*, which are known to be expressed in RMG interneurons (Figure 1–figure supplement 1C). We confirmed RNA-seq data using transcriptional *gfp* reporters of *flp-5* and *flp-21* in RMG (Figure 1F and G). These data predict that *ilc-17*.*1 OE* likely heightens neuropeptidergic signaling to promote the responses to 21% O_2_.

We next sought to determine whether *ilc-17*.*1 OE* in *C. elegans* has any long-term consequences on animals’ physiology. *ilc-17*.*1 OE* animals have much smaller bodies, as well as reduced lifespan, and decreased survival to the exposure of organic peroxide tert-butyl hydroperoxide (tBOOH) (Figure 1–figure supplement 1D; Figure 1H and I). Weakened tBOOH tolerance and shortened longevity of *ilc-17*.*1 OE* animals were suppressed by disrupting *ilcr-1, malt-1, pik-1* or *nfki-1*, encoding the downstream components of ILC-17.1 (Figure 1–figure supplement 1E–J). Meanwhile, *ilc-17*.*1* mutants displayed enhanced tBOOH tolerance (Figure 1I). Disrupting *ilcr-1, malt-1, pik-1* or *nfki-1* led to a similar phenotype (Figure 1–figure supplement 1E–H).

These observations suggest that ILC-17.1 acts through these downstream molecules to modulate longevity and stress response.

Persistent ‘fight-or-flight’ response is known to damage cytoprotective mechanisms, thereby shortening longevity and weakening stress tolerance (De Rosa et al., 2019). Since *ilc-17*.*1 OE* animals are hypersensitive to 21% O_2_ and keep moving fast at 21% O_2_, we wondered whether constant escaping attempts from aversive 21% O_2_ would sacrifice their defense mechanism against stress. However, *npr-1* mutants, which constantly seek to escape from 21% O_2_, were not as tBOOH sensitive as *ilc-17*.*1 OE* animals (Figure 1J). In addition, *ilc-17*.*1 OE* animals, when cultivated for multiple generations at preferred 7% O_2_, still had reduced lifespan and decreased survival to tBOOH exposure (Figure 1–figure supplement 1K; Figure 1K). These data suggest that increased ‘fight-or-flight’ response to 21% O_2_ do not impair cytoprotective mechanisms of *ilc-17*.*1 OE* animals.

### Enhanced ILC-17.1 signaling inhibits NHR-49

To provide molecular insight into ILC-17.1 mediated inhibition of longevity and stress tolerance, we focused on identifying additional components in ILC-17.1 pathway. In the previous study (Flynn et al., 2020), three independent attempts to immunoprecipitate MALT-1 identified NHR-49 (Flynn et al., 2020) (Figure 2A), which belongs to the nuclear hormone receptor family of transcription factors (Antebi, 2006; Lee, Goh, Wong, Klassen, & Taubert, 2016). To confirm their interaction, we performed co-immunoprecipitation (Co-IP) using HA-tagged MALT-1 and GFP-tagged NHR-49. IP of NHR-49-GFP co-precipitated MALT-1-HA; reciprocally, IP of MALT-1-HA brought with it NHR-49-GFP (Figure 2B and C). These results indicate MATL-1 and NHR-49 form a complex.

**Figure 2.**
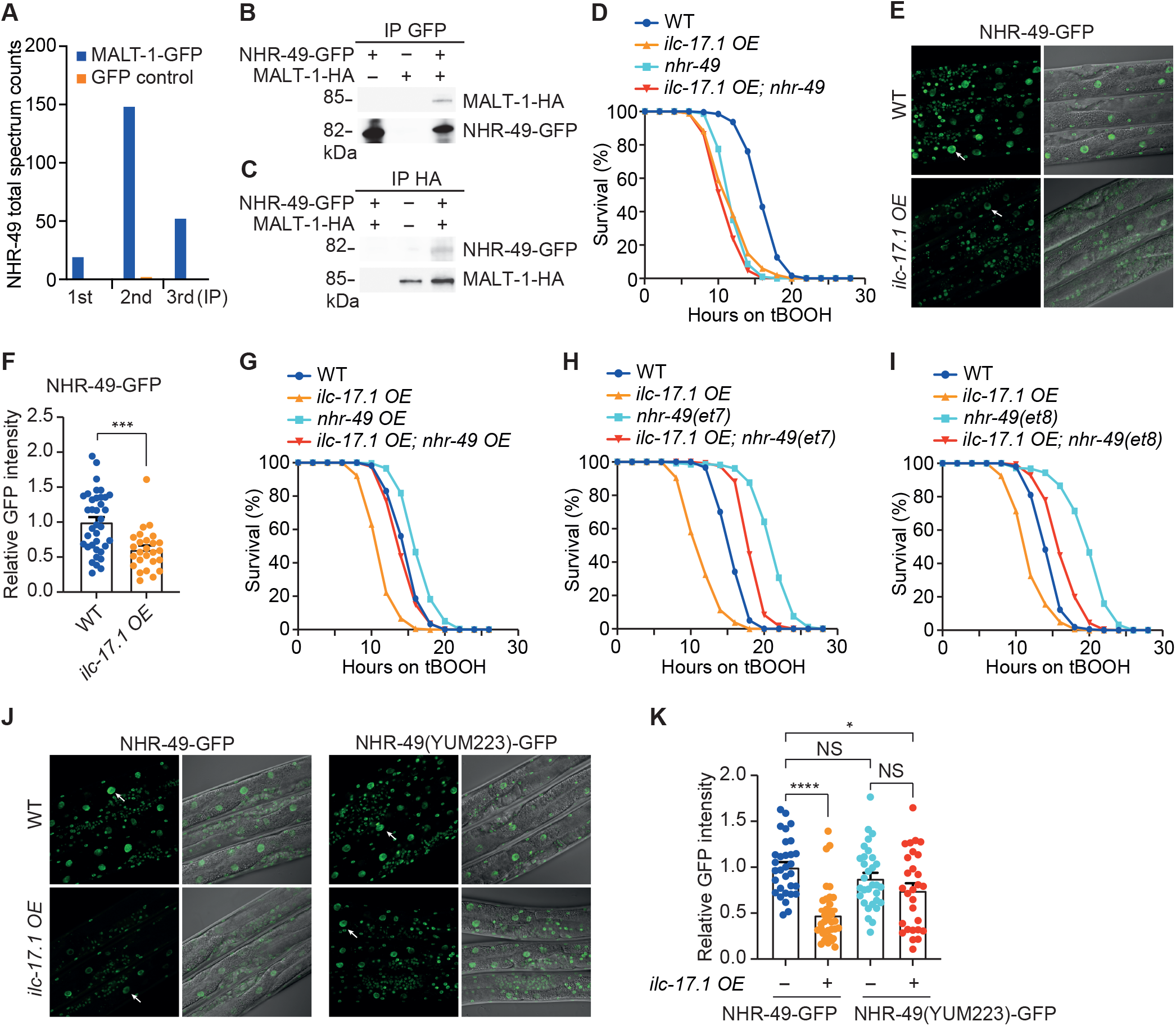
ILC-17.1 overexpression inhibits the activity of nuclear hormone receptor NHR-49. (**A**) Total spectral counts of NHR-49 in three independent immunoprecipitations of MALT-1-GFP with different amount of starting material. GFP expressed from the same promoter was used as control. The data were extracted from (Flynn et al., 2020). (**B**) Immunoprecipitation of endogenously tagged NHR-49-GFP and western blot detection of MALT-1-HA. (**C**) Immunoprecipitation of endogenously tagged MALT-1-HA and western blot detection of NHR-49-GFP. (**D**) Survival plots of WT, *ilc-17*.*1 OE, nhr-49(nr2041)* and *ilc-17*.*1 OE; nhr-49(nr2041)* animals on 7.7mM tBOOH. Numerical values and statistical analyses are in Supplementary File 1. (**E**) Representative images showing endogenously tagged NHR-49-GFP expression in the intestinal nuclei of WT and *ilc-17*.*1 OE* animals. Arrow points to one nucleus in the intestine. (**F**) Quantification of NHR-49-GFP fluorescent intensity in the intestinal nuclei of WT (n=37) and *ilc-17*.*1 OE* animals (n=26). Error bars indicate SEM. ***, *p*<0.001, *t* test. (**G**) Survival curves of WT, *ilc-17*.*1 OE, nhr-49 OE* and *ilc-17*.*1 OE; nhr-49 OE* animals on 7.7mM tBOOH. *nhr-49* overexpression (OE) was from extrachromosomal arrays. Numerical values and statistical analyses are in Supplementary File 1. (**H**) Survival curves of WT, *ilc-17*.*1 OE, nhr-49(et7)* and *ilc-17*.*1 OE; nhr-49(et7)* animals on 7.7mM tBOOH. *et7* is a gain-of-function allele of *nhr-49*. Numerical values and statistical analyses are in Supplementary File 1. (**I**) Survival curves of WT, *ilc-17*.*1 OE, nhr-49(et8)* and *ilc-17*.*1 OE; nhr-49(et8)* animals on 7.7mM tBOOH. *et8* is a gain-of-function allele of *nhr-49*. Numerical values and statistical analyses are in Supplementary File 1. (**J**) NHR-49-GFP (left) and NHR-49(YUM223)-GFP (right) expression in the intestinal nuclei of WT and *ilc-17*.*1 OE* animals. Arrow points to one nucleus in the intestine. *yum223* is a gain-of-function allele of *nhr-49* generated by CRISPR, and is equivalent to *nhr-49(et8)*. (**K**) Quantification of NHR-49-GFP and NHR-49(YUM223)-GFP fluorescent intensity in the intestinal nuclei of WT and *ilc-17*.*1 OE* animals. n=31, 35, 28 and 31 from the left to the right. Error bars indicate SEM. ****, *p*<0.0001, *, *p*<0.05 and NS = not significant. ANOVA with Tukey correction.

NHR-49 has previously been implicated in ageing and stress responses (Dasgupta et al., 2020; Doering et al., 2022; Folick et al., 2015; Goh et al., 2014; Goh et al., 2018; Hu, D’Amora, MacNeil, Walhout, & Kubiseski, 2018; Naim et al., 2021; Wani et al., 2021). It acts as a key regulator of cytoprotective genes during oxidative stress (Goh et al., 2018), and *nhr-49* mutants were hypersensitive to tBOOH (Goh et al., 2014; Goh et al., 2018). Consistent with earlier observations, *nhr-49* null mutants had compromised tBOOH tolerance (Figure 2D). And the *ilc-17*.*1 OE*; *nhr-49* double mutants displayed similar tBOOH sensitivity as the single mutants (Figure 2D), suggesting that in genetic terms ILC-17.1 and NHR-49 act in the same pathway.

Previous work showed that NHR-49-GFP, when overexpressed, was in both nucleus and cytosol (Lee et al., 2016; Ratnappan et al., 2014), suggesting that NHR-49 might shuttle between two cellular compartments. To explore this, we endogenously tagged NHR-49 with GFP. We found that GFP signal was predominantly in the nucleus (Figure 2E). In *ilc-17*.*1 OE* animals, NHR-49-GFP level in the nuclei was dramatically decreased (Figure 2E and F), reflecting that either NHR-49 was redistributed to the cytosol or the overall expression of NHR-49 was decreased in these animals.

However, forced nuclear entry of NHR-49, by tagging endogenously with a SV40 nuclear localization signal (NLS), did not suppress the tBOOH hypersensitivity of *ilc-17*.*1 OE* animals (Figure 2–figure supplement 1A), suggesting that *ilc-17*.*1 OE* likely inhibits the expression of NHR-49. Both RNA-seq and quantitative real-time PCR (qPCR) showed that *nhr-49* transcripts were not reduced in *ilc-17*.*1 OE* animals (Figure 2–figure supplement 1B and C), implying that the inhibition of NHR-49 expression by ILC-17.1 is posttranscriptional.

To address if reduced NHR-49 expression could explain the tBOOH hypersensitivity of *ilc-17*.*1 OE* animals, we overexpressed *nhr-49* (*nhr-49 OE*). This manipulation partially suppressed the phenotype of *ilc-17*.*1 OE* animals (Figure 2G). When *nhr-49* was overexpressed in different tissues, its overproduction in neurons, but not in the other tissues, suppressed the tBOOH hypersensitivity of *ilc-17*.*1 OE* animals (Figure 2–figure supplement 1D–F), suggesting that NHR-49 mainly acts in neurons to regulate tBOOH responses. We next explored if the *gof* alleles of *nhr-49* (*et7* or *et8*), which alters the ligand binding domain of NHR-49 and upregulates its target genes (Lee et al., 2016; Svensk et al., 2013), could rescue the observed phenotype of *ilc-17*.*1 OE* animals. These two alleles of *nhr-49* partially restored tBOOH tolerance to *ilc-17*.*1 OE* animals (Figure 2H and I). To further examine how *nhr-49(gof)* achieved this, we generated a *gof* allele *nhr-49(yum223)* in *nhr-49::gfp* strain using CRISPR/Cas9. This allele is equivalent to *nhr-49(et8)* with a S432F substitution (Svensk et al., 2013). *ilc-17*.*1 OE* failed to repress the expression of NHR-49(YUM223)-GFP (Figure 2J and K), suggesting that gain-of-function mutations uncouple NHR-49 expression from ILC-17.1 activity. Interestingly, both *nhr-49 OE* or *nhr-49(gof)* only partially suppressed tBOOH hypersensitivity of *ilc-17*.*1 OE* animals (Figure 2G–I), and the tBOOH tolerance of *ilc-17*.*1* mutants was incompletely suppressed by disrupting *nhr-49* (Figure 2–figure supplement 1G), suggesting that cellular processes regulated by ILC-17.1 and NHR-49 are not fully overlapping, and they also independently regulate the other biological functions to modulate tBOOH responses.

### ILC-17.1 overexpression inhibits MDT-15

NHR-49 forms a complex with MDT-15, a conserved mediator subunit for transcription coactivation, to collaboratively promote stress tolerance (Goh et al., 2014; Taubert, Van Gilst, Hansen, & Yamamoto, 2006). However, the immunoprecipitation of MATL-1 only pulled down NHR-49, but not MDT-15 (Flynn et al., 2020). Instead, MDT-15 but not NHR-49 was found in the immunoprecipitation of NFKI-1-GFP (Flynn et al., 2020) (Figure 3A), suggesting that NHR-49 and MDT-15 are likely in a larger complex but they independently bind to their own interactors in ILC-17.1 signaling. The interaction between NFKI-1 and MDT-15 was further confirmed by co-IP (Figure 3B and C). Consistent with earlier observations (Goh et al., 2014), *mdt-15* mutants were hypersensitive to tBOOH (Figure 3D). The defect of *ilc-17*.*1 OE* and *mdt-15* on tBOOH sensitivity was not additive (Figure 3D), suggesting that MDT-15 acts in the same genetic pathway as ILC-17.1. This was supported by the observation that increased tBOOH tolerance of *ilc-17*.*1* mutants was suppressed by *mdt-15* loss of function (Figure 3–figure supplement 1A). To further dissect the relationship between ILC-17.1 and MDT-15, we created a *gof* allele *mdt-15*(*yum148*) using CRISPR/Cas9. This allele was equivalent to *mdt-15(et14)*, which has been shown to suppress cold sensitivity of *paqr-2* and activate detoxification responses (Mao et al., 2019; Svensk et al., 2013). *mdt-15*(*yum148*) partially suppressed the tBOOH hypersensitivity of *ilc-17*.*1 OE* animals (Figure 3E), but did not further enhance the tBOOH tolerance of *ilc-17*.*1* mutants (Figure 3–figure supplement 1B), suggesting that *ilc-17*.*1 OE* could inhibit MDT-15. To explore how *ilc-17*.*1 OE* might inhibit MDT-15, we examined the expression and localization of an endogenously tagged MDT-15-GFP. The expression of MDT-15-GFP was significantly decreased in *ilc-17*.*1 OE* (Figure 3–figure supplement 1C and D), and increased *mdt-15* expression partially rescued tBOOH hypersensitivity of *ilc-17*.*1 OE* animals (Figure 3–figure supplement 1E). MDT-15 expression was not affected by *ilc-17*.*1 OE* in animals bearing a *gof* allele *mdt-15(yum596)*, which was also equivalent to *mdt-15(et14)* (Figure 3–figure supplement 1F and G), suggesting that *mdt-15(gof)* might restore tBOOH tolerance to *ilc-17*.*1 OE* animals by maintaining *mdt-15* expression. Similar to NHR-49, *mdt-15(gof)* did not fully suppress the tBOOH hypersensitivity of *ilc-17*.*1 OE* animals (Figure 3E), implying that the molecular functions of ILC-17.1 and MDT-15 are not fully overlapping, and they could act independently to modulate tBOOH responses.

**Figure 3.**
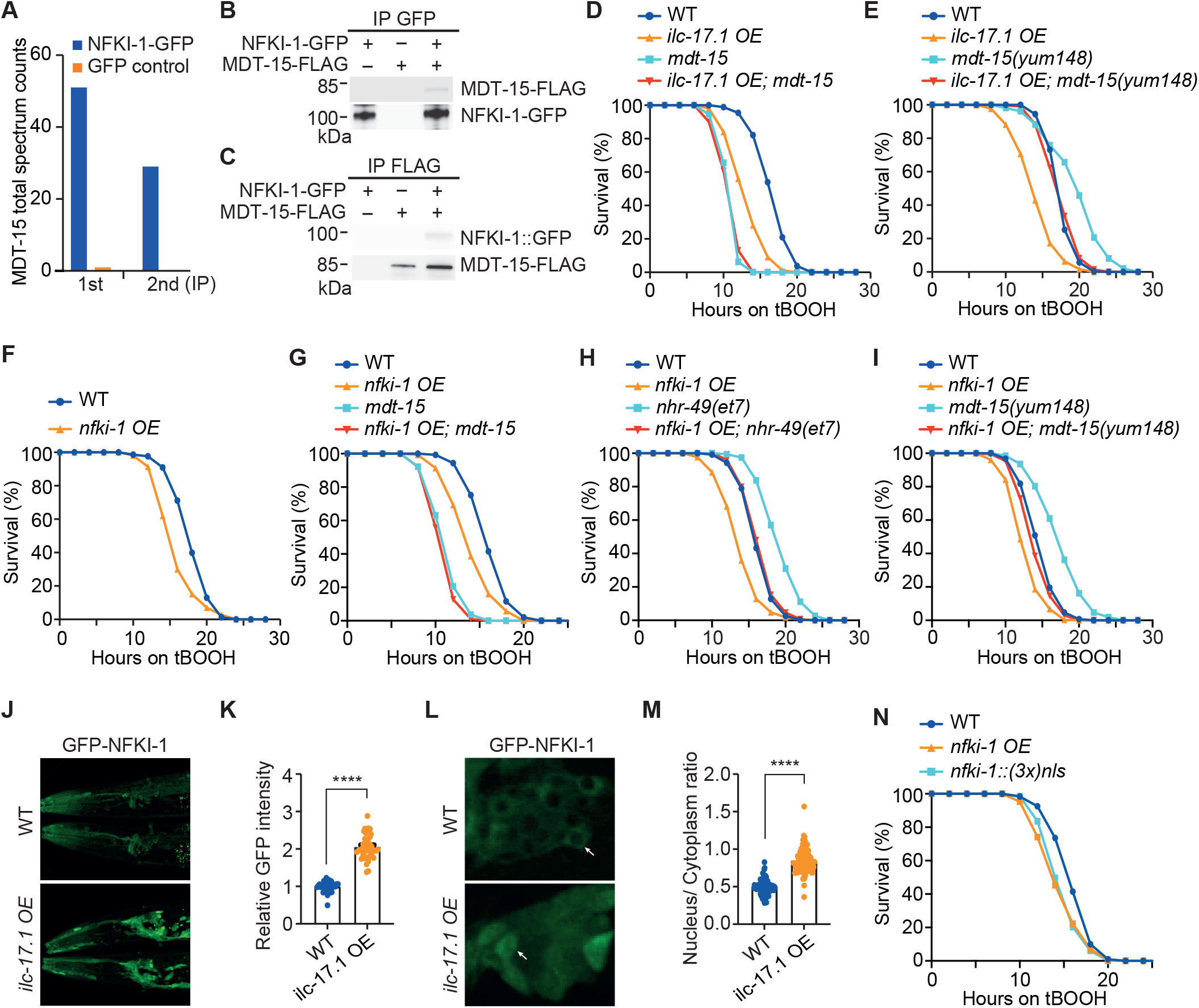
ILC-17.1 negatively regulates the activity of MDT-15. (**A**) Total spectral counts of MDT-15 in two independent immunoprecipitations of NFKI-1-GFP with different amount of starting material. GFP expressed from the same promoter was used as control. The data were extracted from (Flynn et al., 2020). (**B**) Immunoprecipitation of NFKI-1-GFP and western blot detection of MDT-15-FLAG. (**C**) Immunoprecipitation of endogenously tagged MDT-15-FLAG and western blot detection of NFKI-1-GFP. (**D**) Survival curves of WT, *ilc-17*.*1 OE, mdt-15(tm2182)* and *ilc-17*.*1 OE; mdt-15(tm2182)* animals on 7.7mM tBOOH. Numerical values and statistical analyses are in Supplementary File 1. (**E**) Survival curves of WT, *ilc-17*.*1 OE, mdt-15(yum148)* and *ilc-17*.*1 OE; mdt-15(yum148)* animals on 7.7mM tBOOH. *yum148* is a gain-of-function allele of *mdt-15* generated by CRISPR, and is equivalent to *mdt-15(et14)*. Numerical values and statistical analyses are in Supplementary File 1. (**F**) Survival curves of WT and *nfki-1 OE* on 7.7mM tBOOH. *nfki-1 OE* was generated by integrating *nfki-1* extrachromosomal arrays into the genome. Numerical values and statistical analyses are in Supplementary File 1. (**G**) Survival curves of WT, *nfki-1 OE, mdt-15(tm2182)* and *nfki-1 OE*; *mdt-15(tm2182)* animals on 7.7mM tBOOH. Numerical values and statistical analyses are in Supplementary File 1. (**H**) Survival curves of WT, *nfki-1 OE, nhr-49(et7)* and *nfki-1 OE*; *nhr-49(et7)* animals on 7.7mM tBOOH. *et7* is a gain-of-function allele of *nhr-49*. Numerical values and statistical analyses are in Supplementary File 1. **(I)** Survival curves of WT, *nfki-1 OE, mdt-15(yum148)* and *nfki-1 OE*; *mdt-15(yum148)* animals on 7.7mM tBOOH. *yum148* is a gain-of-function allele generated by CRISPR, and is equivalent to *mdt-15(et14)*. Numerical values and statistical analyses are in Supplementary File 1. (**J**) Representative images showing endogenously tagged GFP-NFKI-1 expression in the head region of WT and *ilc-17*.*1 OE* animals. (**K**) Quantification of GFP-NFKI-1 fluorescent intensity in the head of WT (n=35) and *ilc-17*.*1 OE* animals (n=45). Error bars indicate SEM. ****, *p*<0.0001, *t* test. (**L**) Representative images showing endogenously tagged GFP-NFKI-1 expression in the nucleus and cytosol of neurons in WT and *ilc-17*.*1 OE* animals. (**M**) Quantification of GFP-NFKI-1 fluorescent intensity in the nucleus and cytosol of neurons in WT (n=100) and *ilc-17*.*1 OE* animals (n=105). The ratios of fluorescent intensity between nucleus and cytosol were plotted. Error bars indicate SEM. ****, *p*<0.0001, *t* test. (**N**) Survival curves of WT, *nfki-1 OE* and *nfki-1-(3x)nls* animals on 7.7mM tBOOH. Three copies of SV40 nuclear localization signal (3x nls) were inserted before stop codon of *nfki-1* gene on the chromosome. Numerical values and statistical analyses are in Supplementary File 1.

The physical interaction between NFKI-1 and MDT-15 provoked us to investigate if NFKI-1 played a role in the inhibition of MDT-15. We found that reduced MDT-15 expression in *ilc-17*.*1 OE* animals was restored by disrupting *nfki-1* (Figure 3–figure supplement 1H and I), suggesting that ILC-17.1 acts through NFKI-1 to inhibit MDT-15 expression. Interestingly, *nfki-1 OE*, which increased the overall expression of *nfki-1* more than 60-fold (Figure 3–figure supplement 1J), also decreased MDT-15 expression (Figure 3–figure supplement 1K and L), and led to tBOOH hypersensitive (Figure 3F). This phenotype of *nfki-1 OE* was not additive to that of *mdt-15* mutants (Figure 3G), and was partially suppressed by *nhr-49(gof)* or *mdt-15(gof)* alleles (Figure 3H and I). Taken together, these data suggest that *nfki-1 OE* likely inhibits MDT-15 by decreasing MDT-15 expression. We next explored in which tissue NFKI-1 acts to modulate tBOOH responses. *nfki-1 OE* in neurons, but not in the other tissues, was sufficient to render animals hypersensitive to tBOOH (Figure 3–figure supplement 1M–O), suggesting that NFKI-1 acts in neurons.

Similar phenotypes displayed by *ilc-17*.*1 OE* and *nfki-1 OE* animals implied that *ilc-17*.*1 OE* might enhance the expression of NFKI-1. Indeed, *ilc-17*.*1 OE* significantly increased NFKI-1 expression, particularly in the nucleus (Figure 3J–M). Furthermore, tagging NFKI-1 with SV40 NLS at its endogenous locus rendered animals susceptible to tBOOH (Figure 3N), and it partially suppressed the tBOOH tolerance of *ilc-17*.*1* mutants (Figure 3–figure supplement 1P). Collectively, these data suggest that *ilc-17*.*1 OE* inhibits MDT-15 expression by promoting the nuclear levels of NFKI-1.

### ILC-17.1 overexpression represses the expression of NHR-49/MDT-15 target genes

The aforementioned observations suggest that *ilc-17*.*1 OE* in *C. elegans* could alter gene expression by inhibiting the NHR-49/MDT-15 transcription complex. Thus, genes regulated by ILC-17.1 might significantly overlap with NHR-49/MDT-15 targets. To this end, we compared the transcriptional profiles of WT, *ilc-17*.*1 OE, nhr-49(nr2041), nhr-49(et7), nhr-49(et7); ilc-17*.*1 OE, mdt-15(tm2182), mdt-15(yum148)*, and *mdt-15(yum148); ilc-17*.*1 OE* animals (Supplementary File 2). As expected, *ilc-17*.*1* level was dramatically increased in *ilc-17*.*1 OE* animals (Log2 fold change = 10) (Figure 4A). And *ilc-17*.*1 OE* led to a substantial reprograming of gene expression with 401 genes significantly downregulated and 218 genes upregulated (Figure 4A; Supplementary File 2, for adj. *p* <1e-10). Gene Ontology (GO) analysis revealed that cellular processes including defense response to bacterium, and fatty acid biosynthesis were among the most highly down-regulated signatures by *ilc-17*.*1 OE* (Figure 4B, for adj. *p* <1e-10), consistent with the observed phenotypes of *ilc-17*.*1 OE* animals. The down-regulated genes by *ilc-17*.*1 OE* displayed a large degree of overlap with those of *nhr-49* or *mdt-15* mutants (Figure 4C, for adj. *p* <1e-10). Of all the genes down-regulated, 59 of them were shared by all three strains, and 173 genes were jointly repressed in at least two strains (Figure 4C). However, only 9 genes were commonly up-regulated in all three strains (Figure 4D), suggesting that cellular processes enhanced by *ilc-17*.*1 OE* is largely independent of NHR-49/MDT-15. For example, neuronal excitability enhanced by *ilc-17*.*1 OE* was unlikely through the inhibition of NHR-49/MDT-15 (data not shown).

**Figure 4.**
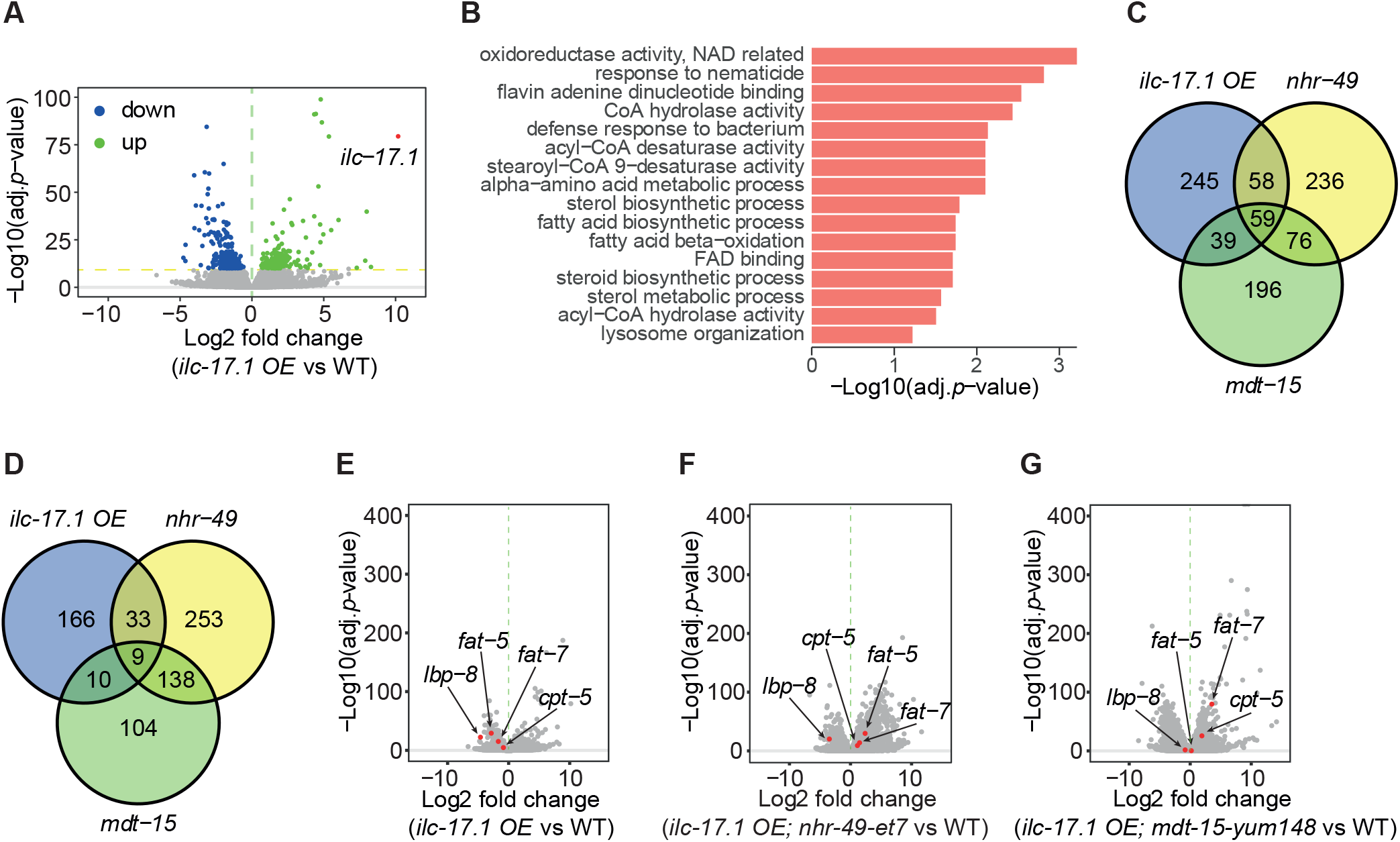
ILC-17.1 overactivation represses transcriptional outputs from NHR-49/MDT-15 complex. (**A**) Volcano plot showing the differentially expressed genes with adjusted *p* value <1e-10 in *ilc-17*.*1 OE* animals. The relative expression of *ilc-17*.*1* is highlighted in red. (**B**) Significantly enriched GO categories of down-regulated genes with adjusted *p* value <1e-10 in *ilc-17*.*1 OE* animals. (**C** and **D**) Venn diagrams displaying the number of significantly down (C) and up (D) regulated genes in *ilc-17*.*1 OE, nhr-49(nr2041)* and *mdt-15(tm2182)* with adjusted *p* value <1e-10. (**E**–**G**) Volcano plots highlighting the expression of NHR-49/MDT-15 target genes, *fat-5, fat-7, cpt-5* and *lbp-8*, in *ilc-17*.*1 OE* (E), in *ilc-17*.*1 OE; nhr-49(et7)* (F) and in *ilc-17*.*1 OE; mdt-15(yum148)* (G) animals relative to wild type.

The *gof* alleles of *nhr-49* and *mdt-15* restored the expression of many down-regulated genes in *ilc-17*.*1 OE* animals (Figure 4–figure supplement 1A and B; Supplementary File 2, sheets 8,9). Among those 401 down-regulated genes, the expression of 55 and 90 genes was increased more than 2-fold in *nhr-49(et7); ilc-17*.*1 OE* and in *mdt-15(yum148); ilc-17*.*1 OE* relative to *ilc-17*.*1 OE* animals (Figure 4–figure supplement 1A and B; Supplementary File 2, sheets 8,9). We also noticed that the expression of 3 genes was further repressed by *nhr-49(et7)* for more than 2-fold (Figure 4–figure supplement 1A; Supplementary File 2, sheets 8,9). Surprisingly, the expression of 51 genes was repressed in *mdt-15(yum148); ilc-17*.*1 OE* animals (Figure 4–figure supplement 1B; Supplementary File 2, sheets 8,9). These observations suggest that ILC-17.1 and NHR-49/MDT-15 commonly regulate the expression of a fraction of genes, but they also act in parallel to modulate transcription. Examining the expression of known NHR-49 and MDT-15 targets, *fat-5, fat-7, lbp-8* and *cpt-5* (Taubert et al., 2006), we found that their expression was all downregulated in *ilc-17*.*1 OE* animals (Figure 4E), and was restored to the various degrees by *nhr-49(et7)* or *mdt-15(yum148)* (Figure 4F and G; Figure 4–figure supplement 1A and B). We validated RNA-seq data by qPCR, confirming that the expression of these genes was significantly decreased in *ilc-17*.*1 OE* animals (Figure 4–figure supplement 1C). The reduced expression was partially restored in *nhr-49(et7); ilc-17*.*1 OE*, or *mdt-15(yum148); ilc-17*.*1 OE* animals (Figure 4–figure supplement 1D and E), which were still lower than those in *nhr-49(et7)* or *mdt-15(yum148)* single mutants. These data are consistent with the partial suppression of *ilc-17*.*1 OE* phenotypes by these *gof* mutants. Therefore, we conclude that NHR-49/MDT-15 complex contributes to the transcriptional outputs in *ilc-17*.*1 OE* animals, but ILC-17.1 and NHR-49/MDT-15 also act independently to regulate gene expression.

### ILC-17.1 overactivation inhibits the lysosome-to-nucleus signaling

Comparing transcriptional profiles of WT and *ilc-17*.*1 OE* animals further revealed that the expression of lysosomal lipase (*lipl*) genes was repressed to various extent by *ilc-17*.*1 OE* (Figure 5A). We validated RNA-seq data using transcriptional *gfp* reporters of *lipl-1, 2, 3* and *5* (Figure 5B and C). Previous observation (O’Rourke & Ruvkun, 2013) together with our RNA-seq analysis showed that *lipl-1* to *lipl-4* were expressed at very low levels under normal conditions. We were surprised to see bright GFP signals from the transcriptional *gfp* reporters of these *lipl* genes (Figure 5B), which were possibly due to the overexpression of these constructs from extrachromosomal arrays. Lysosomal lipases have been shown to modulate longevity and stress resistance (Folick et al., 2015; O’Rourke, Kuballa, Xavier, & Ruvkun, 2013; O’Rourke & Ruvkun, 2013; Ramachandran et al., 2019), and act through NHR-49 in a lysosome-to-nucleus signaling (Folick et al., 2015). To examine if reduced expression of *lipl* genes contributed to the phenotype of *ilc-17*.*1 OE* animals, we simultaneously overexpressed *lipl-1,2,3,4* (*lipl OE*). This manipulation partially suppressed tBOOH susceptibility of *ilc-17*.*1 OE* animals (Figure 5D), and it did not further enhance the tBOOH tolerance of *ilc-17*.*1* mutants (Figure 5–figure supplement 1A). It has been suggested that *lipl OE* promotes the accumulation of oleic acid (OA) and oleoylethanolamide (OEA) (Folick et al., 2015). We assumed that OA and OEA levels might be reduced in *ilc-17*.*1 OE* animals due to decreased *lipl* gene expression, and play a role in regulating tBOOH responses. Indeed, dietary supplementation of OA and KDS-5104, an analogue of OEA (Folick et al., 2015; Fu et al., 2003), partially rescued tBOOH hypersensitivity of *ilc-17*.*1 OE* animals (Figure 5E and F).

**Figure 5.**
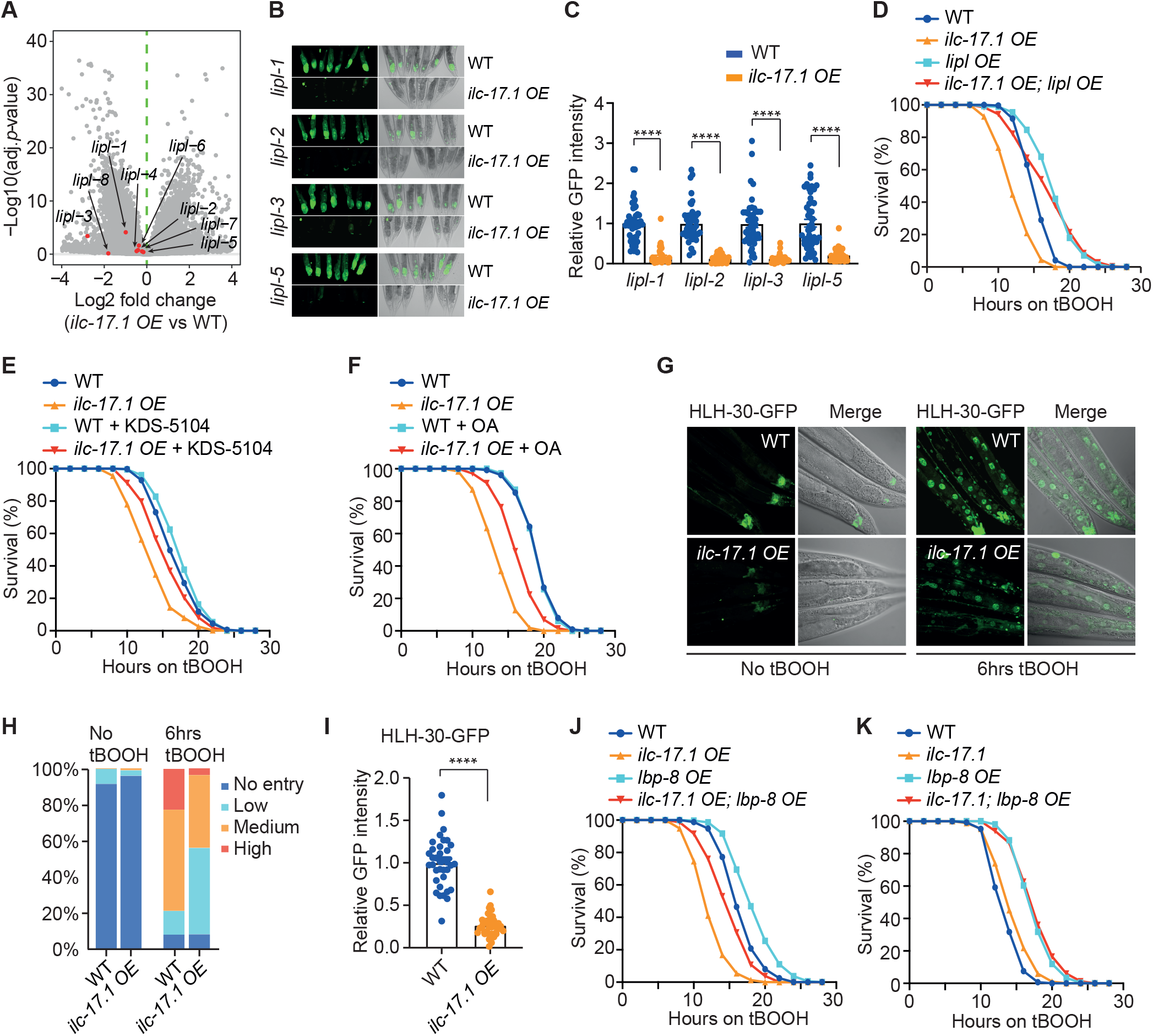
Increased ILC-17.1 signaling inhibits lysosome-to-nucleus signaling. (**A**) Volcano plot highlighting *lipl* gene expression in *ilc-17 OE* animals relative to wild type. (**B**) Representative images showing *gfp* expression driven by the promoters of *lipl-1, lipl-2, lipl-3 and lipl-5* genes in WT and *ilc-17*.*1 OE* animals. (**C**) Quantification of GFP expression driven by *lipl-1, lipl-2, lipl-3* and *lipl-5* promoters in WT and *ilc-17*.*1 OE* animals. n=47, 40, 47, 29, 45, 30, 48, and 32 from the left to the right. Error bars indicate SEM. ****, *p*<0.0001, *t* test. (**D**) Survival curves of WT, *ilc-17*.*1 OE, lipl OE* and *ilc-17*.*1 OE; lipl OE* animals on 7.7mM tBOOH. *lipl OE* is the overexpression of *lipl-1, lipl-2, lipl-3* and *lipl-4* from extrachromosomal arrays. Numerical values and statistical analyses are in Supplementary File 1. (**E**) Survival curves of WT and *ilc-17*.*1 OE* animals with or without oleoylethanolamide (OEA) analogue KDS-5104. Numerical values and statistical analyses are in Supplementary File 1. (**F**) Survival curves of WT and *ilc-17*.*1 OE* animals with or without oleic acid (OA). Numerical values and statistical analyses are in Supplementary File 1. (**G**) Representative images showing the nuclear localization of HLH-30-GFP in WT and *ilc-17*.*1 OE* animals with or without 6 hours of tBOOH exposure. (**H**) Quantification of the nuclear localization of HLH-30-GFP in WT and *ilc-17*.*1 OE* animals with or without 6 hours of tBOOH exposure. n=282, 258, 322 and 263 from the left to the right. (**I**) Quantification of HLH-30-GFP intensity without tBOOH exposure in WT (n=36) and *ilc-17*.*1 OE* animals (n=35). Error bars indicate SEM. ****, *p*<0.0001, *t* test. (**J**) Survival curves of WT, *ilc-17*.*1 OE, lbp-8 OE* and *ilc-17*.*1 OE; lbp-8 OE* animals on 7.7mM tBOOH. *lbp-8 OE* is the overexpression of *lbp-8* from extrachromosomal arrays. Numerical values and statistical analyses are in Supplementary File 1. (**K**) Survival curves of WT, *ilc-17*.*1(tm5218), lbp-8 OE* and *ilc-17*.*1(tm5218); lbp-8 OE* animals on 7.7mM tBOOH. Numerical values and statistical analyses are in Supplementary File 1.

In the RNA-seq analysis we found that the expression of *lipl* genes was not restored in *nhr-49(et7); ilc-17*.*1 OE* or *mdt-15(yum148); ilc-17*.*1 OE* animals (Figure 5–figure supplement 1B and C), suggesting that NHR-49/MDT-15 is not directly required for the expression of *lipl* genes. It has been shown that the transcription factor HLH-30/TFEB is translocated into the nucleus for the induction of *lipl* gene expression under stress (O’Rourke & Ruvkun, 2013). We found that HLH-30 was accumulated in the nucleus after 6 hours of tBOOH exposure, which was inhibited by *ilc-17*.*1 OE* (Figure 5G and H). But forced HLH-30 entry into nucleus failed to restore tBOOH tolerance to *ilc-17*.*1 OE* animals (Figure 5–figure supplement 1D). Further analysis revealed that HLH-30-GFP levels were significantly reduced in *ilc-17*.*1 OE* animals under normal conditions (Figure 5G and I), implying that *ilc-17*.*1 OE* mainly represses the expression of HLH-30.

Lysosomal lipases promote lifespan and stress resistance by activating lipid binding proteins (LBP) (Folick et al., 2015; Savini et al., 2022). RNA-seq analysis showed that the expression of all *lbp* genes, except for *lbp-9*, was decreased in *ilc-17*.*1 OE* animals (Figure 5–figure supplement 1E). Particularly, the NHR-49/MDT-15 target *lbp-8* was one of the most repressed genes by *ilc-17*.*1 OE* (Figure 4E; Figure 5–figure supplement 1E). We confirmed this using a *gfp* reporter of *lbp-8* under its own promoter (Figure 5–figure supplement 1F and G). Increased *lbp-8* expression (*lbp-8 OE*) partially suppressed tBOOH susceptibility of *ilc-17*.*1 OE* animals (Figure 5J). Similar to *lipl OE*, increased tBOOH tolerance by *lbp-8 OE* and *ilc-17*.*1* mutation was not additive (Figure 5K). These observations confirm that *ilc-17*.*1 OE* inhibits lysosomal lipase signaling.

We next explored the possibility that HLH-30/LIPL/LBP acts through NHR-49 to regulate tBOOH responses. *lipl OE* only weakly improved survival of *nhr-49* mutants (Figure 5–figure supplement 1H). Similarly, *lbp-8 OE* had a marginal effect on *nhr-49* mutants (Figure 5–figure supplement 1I), implying that NHR-49 might be one of the downstream effectors of lysosomal lipase signaling. Consistently, the tBOOH tolerance of *hlh-30-(3x)nls* animals was fully suppressed by disrupting *nhr-49* (Figure 5–figure supplement 1J). In addition, RNA-seq data showed that *nhr-49(gof)* failed to restore the reduced expression of *lipl* genes in *ilc-17*.*1 OE* animals (Figure 5–figure supplement 1B). These data confirm the earlier observations that NHR-49 might be downstream of lysosomal lipase pathway.

Tissue specific *lipl OE* revealed that LIPL proteins acted in multiple tissues to promote stress tolerance (Figure 5–figure supplement 1K–M). And *lipl OE* in the intestine displayed the best suppression of tBOOH hypersensitivity of *ilc-17*.*1 OE* animals (Figure 5–figure supplement 1K–M), which was different from the neuronal function of NFKI-1 and NHR-49. This raises the possibility that *ilc-17*.*1 OE* generates a signal from the neurons to modulate *lipl* gene expression in the distal tissues. EGL-21 is a carboxypeptidase for neuropeptide processing. Mutating *egl-21* suppressed tBOOH hypersensitivity of *ilc-17*.*1 OE* animals (Figure 5–figure supplement 1N), suggesting that *ilc-17*.*1 OE* likely enhances neuropeptide secretion from the nervous system (Figure 1–figure supplement 1C) to remotely inhibit the expression of *lipl* genes in the other tissues.

### Increased ILC-17.1 signaling represses CBP/p300 activity

One predominant interactor of NFKI-1-GFP obtained in the previous immunoprecipitations was CBP-1 (Flynn et al., 2020) (Figure 6A), the *C. elegans* ortholog of human CBP/p300 histone acetyltransferase (Arany, Sellers, Livingston, & Eckner, 1994). Depleting *cbp-1* by feeding RNAi attenuated tBOOH tolerance (Figure 6B). A *gof* allele *cbp-1(ku258)* (Eastburn & Han, 2005) rendered animals resistant to tBOOH (Figure 6C and D), and partially restored tBOOH tolerance to *ilc-17*.*1 OE* or *nfki-1 OE* animals (Figure 6C and D), suggesting that either increased ILC-17.1 signaling inhibits CBP-1 or they act in parallel. We further explored this by examining the possible interaction between NHR-49/MDT-15 and CBP-1. Mutating *nhr-49* or *mdt-15* did not further enhance tBOOH hypersensitivity of *cbp-1* inactivation (Figure 6E). And the increased tBOOH tolerance of *nhr-49(gof)* or *mdt-15(gof)* mutants was partially abolished by *cbp-1* RNAi (Figure 6F). These data suggest that CBP-1 is required for the activation or the expression of NHR-49/MDT-15. Interestingly, the expression of *nhr-49* and *mdt-15* was not altered by *cbp-1* RNAi (Figure 6–figure supplement 1A–D), and the *cbp-1* level was not changed in the loss of function or gain of function mutants of *nhr-49* and *mdt-15* (Figure 6–figure supplement 1E–H).

**Figure 6.**
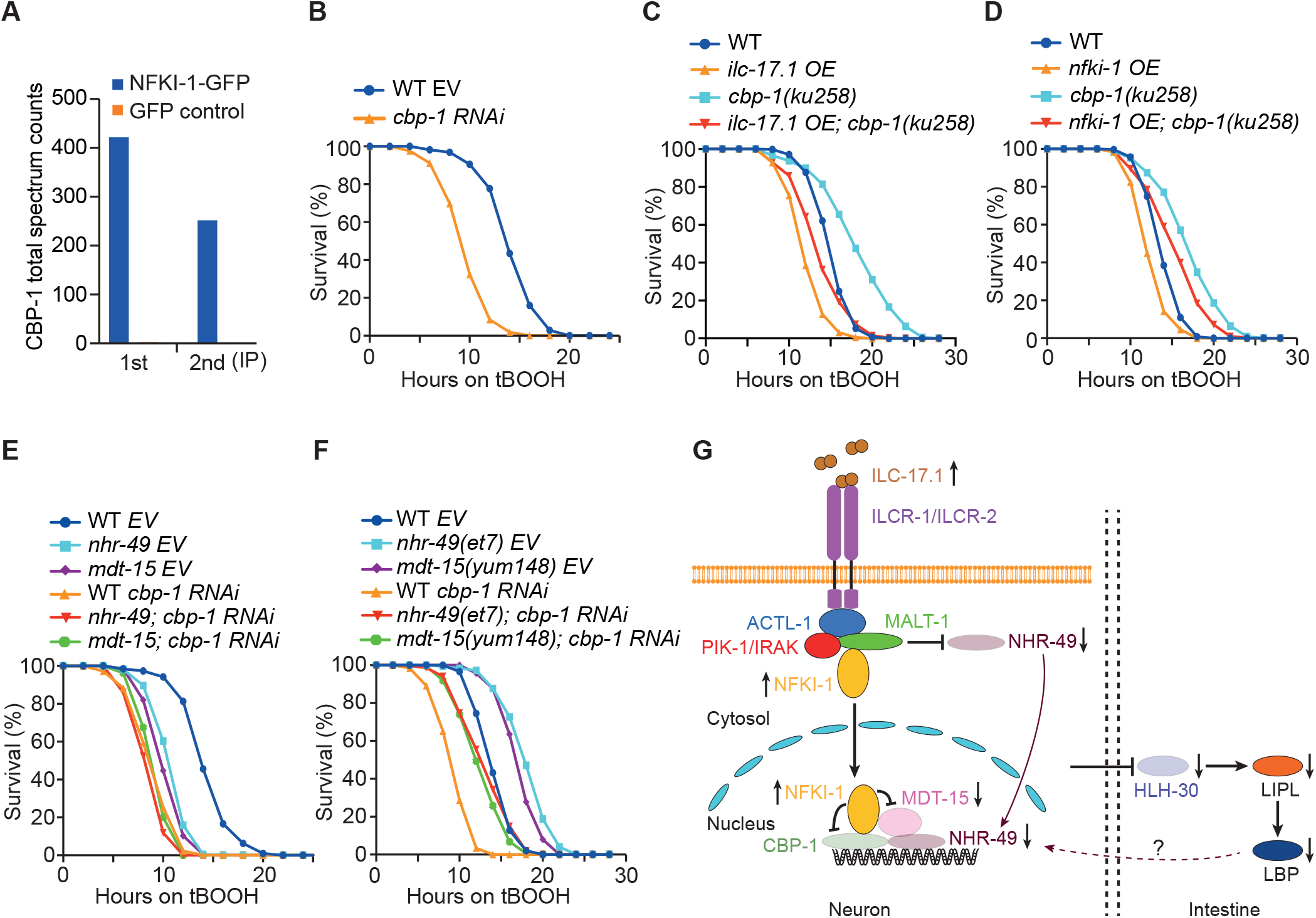
ILC-17.1 signaling inhibits CBP/p300. (**A**) Total spectral counts of CBP-1 in two independent immunoprecipitations of NFKI-1-GFP with different amount of starting material. GFP expressed from the same promoter was used as control. The data were extracted from (Flynn et al., 2020). (**B**) Survival curves of WT animals treated with either RNAi empty vector (EV) L4440 (control) or a RNAi plasmid targeting *cbp-1. cbp-1* RNAi culture was diluted (1:20) by the control before seeding on the RNAi plates. Numerical values and statistical analyses are in Supplementary File 1. (**C**) Survival curves of WT, *ilc-17*.*1 OE, cbp-1(ku258)* and *ilc-17*.*1 OE*; *cbp-1(ku258)* animals on 7.7mM tBOOH. *ku258* is a gain-of-function allele of *cbp-1*. Numerical values and statistical analyses are in Supplementary File 1. (**D**) Survival curves of WT, *nfki-1 OE, cbp-1(ku258)* and *nfki-1 OE*; *cbp-1(ku258)* animals on 7.7mM tBOOH. Numerical values and statistical analyses are in Supplementary File 1. (**E**) Survival curves of WT, *nhr-49(nr2041)* and *mdt-15(tm2182)* animals treated with either the RNAi empty vector (EV) L4440 (control) or a RNAi plasmid targeting *cbp-1. cbp-1* RNAi culture was diluted (1:20) by the control before seeding on the RNAi plates. Numerical values and statistical analyses are in Supplementary File 1. (**F**) Survival curves of WT, *nhr-49(et7)* and *mdt-15(yum148)* animals treated with either the RNAi empty vector (EV) L4440 (control) or a RNAi plasmid targeting *cbp-1. cbp-1* RNAi culture was diluted (1:20) by the control before seeding on the RNAi plates. Numerical values and statistical analyses are in Supplementary File 1. (**G**) Model for the inhibition of a transcriptional network by enhanced ILC-17.1 signaling. *ilc-17*.*1 OE* increases the nuclear expression of NFKI-1, but decreases the expression of NHR-49 and its coactivator MDT-15, and possibly inhibits CBP-1. These components act in neurons to modulate tBOOH responses. In addition, *ilc-17*.*1 OE* remotely inhibits the expression of HLH-30/TFEB, thereby reducing the expression of *lipl* genes in the intestine and damaging the lysosomal lipase signaling in the intestine. The question remains if reduced lysosomal lipase signaling in the intestine leads to compromised activation of NHR-49 in the neurons.

These observations exclude the possible transcriptional regulation between NHR-49/MDT-15 and CBP-1.

Collectively, we described an inhibitory network on transcription when ILC-17.1 is overactivated. *ilc-17*.*1* OE initiates a cascade of reactions to minimize the transcriptional outputs from NHR-49 (Figure 6G), including the reduced expression of NHR-49 and its coactivator MDT-15, the repressed expression of HLH-30 and its target genes, and the inhibition of CBP-1. These responses collaboratively undermine host fitness and stress tolerance of *ilc-17*.*1 OE* animals.

## Discussion

Aberrant activation of IL-17 signaling not only sensitizes the nociceptors in the progression of mechanical hyperalgesia (Ebbinghaus et al., 2017; Luo et al., 2019; Richter et al., 2012), but also underlies the pathogenesis of several autoimmune diseases including multiple sclerosis and rheumatoid arthritis (Gaffen, 2009; Lubberts, 2008; Pappu et al., 2010). We found that increased ILC-17.1 signaling in *C. elegans* not only increases the excitability of O_2_ sensing circuit to aversive 21% O_2_, but also shortens lifespan and weakens stress tolerance. It suggests that enhanced responses to 21% O_2_ could come with the cost of damaging general fitness and cytoprotective mechanisms in *C. elegans*. However, O_2_ sensitive strain *npr-1* is not as susceptible to tBOOH as *ilc-17*.*1 OE* animals, and cultivation at 7% O_2_ fails to rescue shortened lifespan and tBOOH hypersensitivity of *ilc-17*.*1 OE* animals. Thus, compromised longevity and stress tolerance of *ilc-17*.*1 OE* animals are unlikely the consequences of continuous attempts to escape 21% O_2_. But we cannot exclude the possibility that *ilc-17*.*1 OE* enhances the responsiveness to the other noxious stimuli, which jointly impair the defensive mechanisms towards threats.

As a major pro-inflammatory cytokine with critical roles in host defense, IL-17 reprograms gene expression by activating the downstream transcription complex NF-κB (Gaffen, 2009). Interestingly, *C. elegans* does not seem to have canonical NF-κB like transcription factors (Irazoqui et al., 2010). This raised the question how ILC-17.1 signaling in *C. elegans* modulates transcriptional outputs. Our analysis identified NHR-49 as a key element in the transcriptional network downstream of ILC-17.1 signaling (Figure 6G). First, NHR-49 protein level is decreased when *ilc-17*.*1* is overexpressed. Second, *ilc-17*.*1 OE* inhibits the expression of HLH-30/TFEB transcription factor, which leads to the low expression of *lipl* genes and decreases the lysosomal signaling to NHR-49. Third, *ilc-17*.*1 OE* enhances NFKI-1 expression and promotes its nuclear entry, where NFKI-1 inhibits the NHR-49 coactivator MDT-15 and CBP-1. Furthermore, the expression of MDT-15 is reduced in *ilc-17*.*1 OE* animals. Collectively, increased ILC-17.1 signaling triggers an inhibitory network to constrain the activity of NHR-49. We also noticed that *ilc-17*.*1 OE* phenotypes were only partially suppressed by increased NHR-49/MDT-15 signaling, elevated *lipl* expression or enhanced CBP-1 activity, suggesting that ILC-17.1 acts partly in parallel to these molecules to modulate tBOOH responses. Consistently, RNA-seq analysis showed that down-regulated genes in *ilc-17*.*1 OE* animals displayed an extensive overlapping with those in *nhr-49/mdt-15* mutants, but the up-regulated genes overlapped marginally. In addition, the expression of many down-regulated genes was not restored to *ilc-17*.*1 OE* animals by the gain of function mutations of *nhr-49* and *mdt-15*. These data confirmed that the molecular functions of ILC-17.1 and NHR-49/MDT-15 are not fully overlapping, and they also act independently to modulate cellular processes.

Tissue specific expression experiments suggest that NFKI-1 and NHR-49 act in neurons, while lysosomal lipases function mainly in the intestine to regulate tBOOH responses. Interestingly, the expression of *lipl* genes in the intestine is dramatically reduced in *ilc-17*.*1 OE* animals. These observations suggest that ILC-17.1 signaling regulates a neuron-to-intestine signal to modulate *lipl* gene expression in the distal tissues. The suppression of *ilc-17*.*1 OE* phenotypes by *egl-21* implies that enhanced neuropeptidergic signaling is probably involved in the remote inhibition of *lipl* gene expression. Recently, it was reported that *lipl OE* enhanced LBP secretion from the intestine, which was subsequently transported into neurons to regulate the activity of nuclear hormone receptors, establishing a signaling pathway from the periphery tissue to the neurons (Savini et al., 2022). Based on this, it is intriguing to speculate that reduced expression of *lipl* and *lbp* in the intestine of *ilc-17*.*1 OE* animals might insufficiently transmit the intestinal signal to the neurons, and fail to activate NHR-49. Additional experiments are required to fully delineate the ILC-17.1-OE-evoked information flow from the neurons to the intestine, and subsequently back again into the neurons. Nevertheless, our work begins to elucidate the molecular mechanism underlying the long-term consequences of aberrant ILC-17.1 signaling.

## Materials and Methods

### Strains

*C. elegans* were maintained under standard laboratory conditions (Brenner, 1974), unless otherwise stated. Strains used in this study were listed in Supplementary File 3.

### Molecular biology

Plasmids were constructed using multi-site Gateway system (Thermofisher). Promoters used in this study include: *ilc-17*.*1* (6k), *nhr-49* (3.2k), *mdt-15* (3k), *lipl-1* (2.9k), *lipl-2* (1.9k), *lipl-3* (2.9k), *lipl-4* (3k), *lipl-5* (1.5k), and *lbp-8* (2.8k). The transcriptional *gfp* reporter strains of *flp-5* and *flp-21* were obtained from CGC. To generate the reporter and overexpression strains of *lipl* genes, polycistronic constructs with their promoters at position one, their cDNAs at position two and *sl2::gfp* sequence at position 3 were assembled to the destination vector with *unc-54 3’UTR* by LR reaction. For the rescue experiments, cDNA or genomic sequence of each gene was amplified by Phusion High-Fidelity PCR kit (New England BioLabs), cloned to pDonor 221, and assembled to position two of the destination vector with different promoters at position one. Detailed primer information is available in Supplementary File 4.

Transgenic animals were generated by injecting the expression vectors together with 50ng/μl coelomocyte marker *Punc-122::mcherry* or *Punc-122::gfp. lipl-1, lipl-2, lipl-3* and *lipl-4* expression vectors were mixed together at 20ng/μl of each for the injection. The other expression vectors were injected at 50ng/μl.

### Behavioral assays

Aggregation and locomotion assays were performed as previously described (Busch et al., 2012; Laurent et al., 2015). In all the assays, L4 animals of each genotype were picked to fresh plates 24 hours before analysis, and day-one adults were used for assays. For bordering and aggregation, assay plates were seeded with a 1cm diameter bacterial lawn 48 hours earlier. 60 adult animals were transferred to the assay plates, and scored 2 hours after picking. Care was taken not to vibrate the assay plates during the whole assay period. Aggregation was counted when 3 or more animals had body contacts. To assay locomotion, assay plates were seeded with OP50 16 hours before analysis. Bacterial lawn border on the assay plates was removed by a PDMS stamp prior to the assays. 25 animals were picked to the assay plates, and sealed with microfluidic chambers. 7% and 21% O_2_ in 50ml syringes were delivered to the chamber at a flow rate of 3ml/min using PHD 2000 Infusion syringe pump (Harvard Apparatus). Telfon valves (AutoMate Scientific), operated by ValveBank Perfusion Controller, was used to rapidly switch between 7% and 21% O_2_. Videos were recorded at 2 frames/second for 2 minutes at each O_2_ concentration using a FLIR camera mounted on a Zeiss Stemi 508 microscope, and analyzed using a custom-designed MatLab program (https://github.com/wormtracker/zentracker).

### CRISPR-Cas9 genome editing

CRISPR-Cas9 mediated genome editing was performed as previously described (Dokshin, Ghanta, Piscopo, & Mello, 2018; Ghanta & Mello, 2020). Detailed guide, donor and primer information were listed in Supplementary File 5. To disrupt genes including *nfki-1(yum123), nfki-1(yum127)* and *malt-1(yum139)*, we knocked-in three stop codons in different frames and simultaneously removed 16 bases (Supplementary File 5). The insertion templates were single strand DNA oligo (ssODN), containing 35 bases of homology arms (italic in Supplementary File 5) flanking the PAM site, three stop codons (red) and a unique restriction site (green) for genotyping. To create the gain of function mutations of *nhr-49* and *mdt-15* including *yum223, yum140, yum144, yum148*, and *yum596*, PAM sites adjacent to the codon of interest were selected. The ssODN repair template contained 35-base homology arms (italic) with one targeting outside of the guide sequence (blue) and the other located outside of the codon of interest (bold) (Supplementary File 5). All the codons between two homology arms on ssODN were mutated to synonymous codons (lower case) so that ssODN was not targeted by CRISPR/Cas9. Short epitope tags and nuclear localization sequence (yellow) were inserted in a similar fashion. To insert large fragments such as GFP, DNA donor template was PCR-amplified with primers that contain homology arms flanking the stop codon, and purified with AMPure XP beads (A63880, Beckman Coulter). The injection mix was prepared by mixing 0.5μl of Cas9 protein (IDT) with 5μl of 0.4μg/μl tracrRNA (IDT) and 2.8μl of 0.4μg/μl crRNA (IDT). The mix was incubated at 37°C for at least 10 minutes before adding 2.2μl of 1μg/μl ssODN and 2μl of 600μg/μl *rol-6* co-injection marker for gene disruption and point mutatons, or adding 1μl of 500μg/μl dsDNA template and 2μl of 600μg/μl *rol-6* co-injection for large insertions. Nuclease free water was used to bring the final volume to 20μl. The injection mixture was centrifuged for at least 2 minutes before use.

### Oxidative stress assays

In all assays, at least two independent experiments were performed at different days, and each experiment contained at least 3 technical replicates as shown in Supplementary File 1. If the data of two experiments were not consistent, a third assay was included. L4 animals of each genotype were picked and grown overnight on NGM plates. Meanwhile, the assay plates containing tert-butyl hydroperoxide (tBOOH) (Sigma, 8140060005) were prepared by adding the chemical to NGM agar medium to the final concentration of 7.7mM just before pouring plates, and dried overnight. In the next day, synchronized day-one adults were transferred to the tBOOH containing plates, and scored every two hours.

### RNA preparation

RNA samples for RNA-seq and qPCR were prepared as described (Flynn et al., 2020). Animals at L4 stage were collected, and 6 independent samples were prepared for each genotype. Animals were washed 3 times in M9 buffer, and frozen in liquid nitrogen after M9 was removed. Samples were disrupted using Bullet blender (Next Advance) in the presence of 1ml of Qiazol Lysis Reagent with 500μl 0.5mm Zirconia beads (BioSpec) at 4°C. RNA was extracted with RNeasy Plus Universal Mini Kit from Qiagen. Library preparation and RNA sequencing was performed at Novogene.

### Longevity assay

Lifespan experiment was performed as described, with minor modifications (Wilkinson, Taylor, & Dillin, 2012). All assays were performed at 25°C. Synchronized day-one adult animals were divided into five fresh NGM plates, which was used as day 0. Animals were transferred to fresh plates every other day until day 8. Animals were scored every day, and counted as dead if they did not respond to both anterior and posterior prodding. To assay longevity at 7% O_2_, worms were maintained and assayed at 7% O_2_ in an O_2_ chamber (Don Whitley M85 workstation).

### Lipid feeding

The assay was performed as previously described (Folick et al., 2015). KDS-5104 (Cayman Chemical) stock solution was prepared in DMSO to the final concentration of 50mM. The stock was diluted in the dilution buffer (3g NaCl, 1ml 1M CaCl_2_, 1ml MgSO_4_ and 25ml 1M KPO_4_ per liter) to 100μM. 85μl of this was added to the 3.5cm plate, which contained 5ml of NGM medium and was seeded with 100μl concentrated OP50. For oleic acid (OA) feeding, OA was dissolved in ethanol to 600mM, and added to melted NGM agar to the final concentration of 600μM just before pouring plates. The assay was performed in the same way as KDS-5104.

### Quantitative RT-PCR

cDNA was synthesized using Transcriptor High Fidelity cDNA synthesis kit (Sigma, 5091284001). Quantitative RT-PCR was performed using FastStart Universal SYBR Green Master (Rox) (Sigma, 4913850001) on a Bio-Rad CFX Connect Real-Time PCR detection system. *act-1* and *tba-1* were used as internal controls for normalization. Each qPCR quantification included three independent experiments with 9 technical replicates in total. Primers used for qPCR were listed in Supplementary File 4.

### RNA interference

RNAi was performed as previously described (Ahringer, 2006). *E. coli* HT115 (DE3) strain harboring *cbp-1* RNAi vector was obtained from the Ahringer RNAi library. The empty vector L4440 was used as the control. *cbp-1* RNAi culture was diluted (1:20) by the control before seeding on the plates. N2 adult animals were picked to *cbp-1* RNAi plates and laid eggs for 3 hours before removed. Eggs were allowed to develop to day-one adults, which were then used for the assays.

### RNA-seq analysis

Reads were aligned to the *C. elegans* reference genome, WBcel235.103, using the STAR aligner (https://www.ncbi.nlm.nih.gov/pmc/articles/PMC3530905/). The genomic index was generated using flags --sjdbOverhang 150 -- genomeSAindexNbases 10. Counts for each gene were summarized using featureCounts (https://pubmed.ncbi.nlm.nih.gov/24227677/) and then analyzed using R and DESeq2 (https://genomebiology.biomedcentral.com/articles/10.1186/s13059-014-0550-8). Samples were compared qualitatively using a clustering over variance stabilized counts, with replicates clustering together. Pairwise differential expression was performed. Venn diagrams were plotted using the ggvenn package. The GO analysis was performed using the enrichR package, including differentially expressed genes with adjusted *p*-value < 1e-10 and negative fold change.

### Co-immunoprecipitation for Western blot

The worms were grown on 9cm plates at 20°C, and harvested using M9 buffer before starvation. Worm pellet was washed 3 times with M9 and once with ice-cold lysis buffer (50mM HEPES pH7.4, 1mM MgCl_2_, 150mM KCl, 1mM EGTA, 10% glycerol, 0.05% NP40, 1mM phenylmethyl sulphonyl fluoride, 0.1mM DTT, and complete EDTA-free protease inhibitor). Pellets were lysed using Bullet blender (Next Advance) at 4°C. The debris was removed by spinning at 16,000 x g at 4°C for 30 minutes. The supernatant was incubated with GFP-Trap beads (gta-20, ChromoTek), anti-HA agarose (A2095, Sigma) or anti-Flag M2 affinity gel (A2220, Sigma) for 2 hours at 4°C. The agarose beads were subsequently rinsed 3 times with washing buffer (50mM HEPES at pH7.4, 1mM MgCl_2_, 150mM KCl, 1mM EGTA, 10% glycerol, 0.05% NP40), mixed with 30μl of sample buffer, and heated at 95°C for 5 minutes.

1μl was loaded on the gel for the detection of directly immunoprecipitated proteins to avoid overexposure, and the rest of the samples was loaded on TGX Stain-Free protein gels for the detection of co-immunoprecipitated protein. Primary antibodies used in this study include anti-gfp (1:2000, abcam; ab290), anti-HA (1:2000, abcam; ab18181) or anti-flag (1:2000, Sigma; A8592). The chemiluminescence signal was detected using the Odyssey FC imaging systems. Uncropped images can be found in the supporting file.

### Microscopy

Animals were mounted on 2% agarose pads in M9 buffer containing 50mM sodium azide (NaN_3_). Fluorescent images were collected using a Nikon A1 scanning confocal microscope with Nikon NIS elements software. To quantify the *gfp* signal from *lipl* and *lbp-8* constructs, Z-stack images were acquired using synchronized day-one adult animals. To examine the subcellular localization and the expression of NHR-49-GFP, MDT-15-GFP and NFKI-1-GFP, animals at L4 stage were aligned, and Z-stack images were collected and analyzed. Images were taken within 5 minutes after animals were mounted on the agarose pad. We chose to quantify NHR-49-GFP and MDT-15-GFP signals of intestinal nuclei because of their big size, clear boundaries and better signals. In neurons, the NHR-49-GFP and MDT-15-GFP signals were relatively weak, clustered together, and constantly interfered by signals from pharyngal muscle and hypodermis. To quantify the nucleus localization of HLH-30-GFP, synchronized L4 animals were transferred to NGM plates containing 7.7mM tBOOH and grown for 6 hours. Animals were immediately immobilized in 50mM NaN_3_ on the agarose pads. Mounted animals were imaged within 5 minutes. The nucleus localization of HLH-30-GFP in the intestine cells was quantified. To examine the induction of *ilc-17*.*1* expression by *E. faecalis*, the transcriptional mCherry reporter under *ilc-17*.*1* promoter was used (*Pilc-17*.*1::ilc-17*.*1::sl2::mCherry*). *E. faecalis* were grown in brain-heart infusion (BHI) liquid media at 37°C overnight, seeded onto BHI media plates, and incubated at 37 °C for 24 hours. The OP50 seeded on the BHI plates were used as control. The reporter strain at L4 stage was cleaned before the assay started. The images were collected after 4, 8, 12 or 24 hours of infection. All images were processed using Fiji ImageJ.

### Calcium imaging

Transgenic L4 animals expressing calcium sensor YC2.60 in RMG interneurons were picked 24 hours before imaging. Day-one animals were glued onto a 2% agarose pad using Dermabond tissue adhesive on a glass slide, and the nose and tail were exposed in a mix of OP50 and M9 buffer. Animals were sealed in a microfluidic chamber, and O_2_ was delivered into the chamber using the same set-up as that in the behavior analysis. Calcium imaging was performed on an inverted microscope (Nikon) with a 40x lens, and videos were recorded at 2 frames/sec with 100ms exposure time using MetaMorph acquisition software (Molecular Devices). The excitation light was passed through a cyan excitation filter (Chroma), and the emission light was separated for CFP (460–495nm) and YFP (525–580nm). Imaging data were analyzed using a custom written Matlab program (https://github.com/neuronanalyser/neuronanalyser).

### Resource availability

Further information and requests for resources and reagents should be directed to and will be fulfilled by the lead contact Changchun Chen.

Plasmids and strains generated in this study are available upon request from the lead contact.

## Data availability

Raw RNA-seq data have been deposited in ArrayExpress with accession number E-MTAB-11701.

## Acknowledgements

We thank the Caenorhabditis Genetics Center (funded by NIH Office of Research Infrastructure Programs P40 OD010440) and the National BioResources Project Japan for strains. The computations were enabled by resources in project snic2022-5-18 provided by the Swedish National Infrastructure for Computing (SNIC) at UPPMAX, partially funded by the Swedish Research Council through grant agreement no. 2018-05973.

## Additional information Funding

This work was supported by a Swedish Research Council (Vetenskapsrådet, 2021-06602) to JH, and an ERC starting grant (802653), a Swedish Research Council starting grant (2018-02216) and Knut and Alice Wallenberg Foundation to CC.

## Author contributions

Q.L., I.V., J.H. and C.C. designed experiments. Q.L., I.V., L.Z. and L.N. performed the experiments. J.H. did the RNA-seq data analysis. Q.L., I.V., L.Z., L.N., J.H. and C.C. analyzed the data; Q.L, J.H. and C.C. wrote the paper.

## Supporting information

### Supplementary Figure legend

**Figure 1–figure supplement 1.**
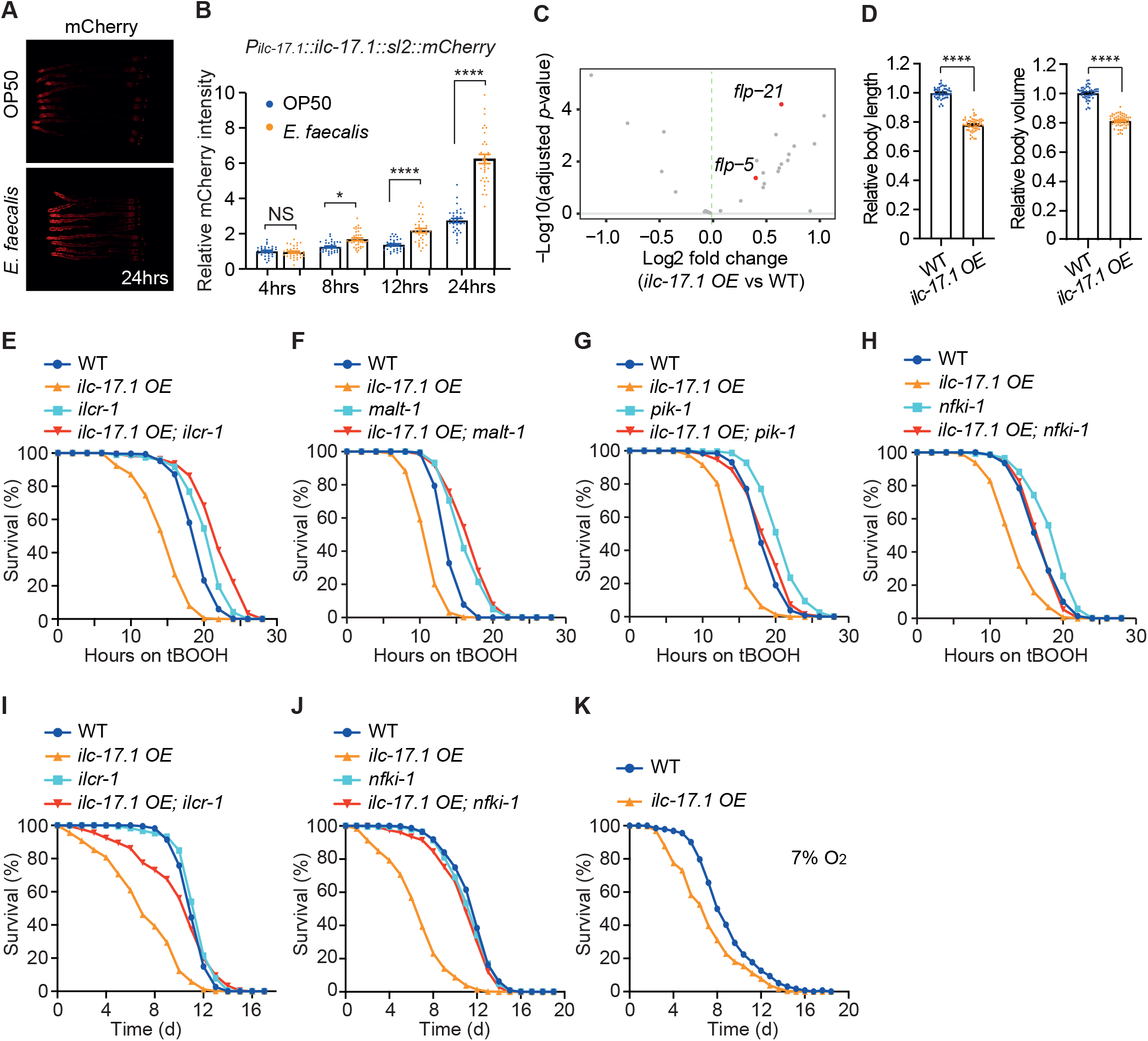
Disrupting downstream components of ILC-17.1 signaling blocks the effects of *ilc-17*.*1 OE*. (**A**) Representative images showing the induction of mCherry expression under the *ilc-17*.*1* promoter by 24-hour of *E. faecalis* exposure. Stage matched animals fed with OP50 were used as the control. (**B**) Quantification of mCherry fluorescent intensity after 4, 8, 12 and 24 hours of *E. faecalis* exposure. The fluorescent intensity of animals from OP50 plates at time 4hrs was arbitrarily set to 1. n=30, 34, 34, 37, 32, 35, 33 and 33 from the left to right. Error bars indicate SEM. NS=not significant, *, *p*<0.05, ****, *p*<0.0001, ANOVA with Tukey correction. (**C**) Volcano plot showing all FMRFamide-like neuropeptide genes in *ilc-17*.*1 OE* animals relative to wild type, and highlighting *flp-5* and *flp-21* that are expressed in RMG interneurons. (**D**) Body length (left) and body volume (right) of WT (n=51) and *ilc-17*.*1 OE* animals (n=55). Error bars indicate SEM. ****, *p* < 0.0001, *t* test. (**E**) Survival curves of WT, *ilc-17*.*1 OE, ilcr-1(tm5866)* and *ilc-17*.*1 OE; ilcr-1(tm5866)* animals on 7.7mM tBOOH. Numerical values and statistical analyses are in Supplementary File 1. (**F**) Survival curves of WT, *ilc-17*.*1 OE, malt-1(yum139)* and *ilc-17*.*1 OE; malt-1(yum139)* animals on 7.7mM tBOOH. *yum139* is a null allele of *malt-1* generated by CRISPR. Numerical values and statistical analyses are in Supplementary File 1. (**G**) Survival curves of WT, *ilc-17*.*1 OE, pik-1(tm2167)* and *ilc-17*.*1 OE; pik-1(tm2167)* animals on 7.7mM tBOOH. Numerical values and statistical analyses are in Supplementary File 1. (**H**) Survival curves of WT, *ilc-17*.*1 OE, nfki-1(yum123)* and *ilc-17*.*1 OE; nfki-1(yum127)* animals on 7.7mM tBOOH. Both *yum123* and *yum127* are null alleles of *nfki-1* generated by CRISPR. *ilc-17*.*1* extrachromosomal array was integrated into a locus tightly linked to *nfki-1* gene. Thus, two different alleles were used. Numerical values and statistical analyses are in Supplementary File 1. (**I**) Lifespan survival curves of WT, *ilc-17*.*1 OE, ilcr-1(tm5866)* and *ilc-17*.*1 OE; ilcr-1(tm5866)*. Numerical values and statistical analyses are in Supplementary File 1. (**J**) Lifespan survival curves of WT, *ilc-17*.*1 OE, nfki-1(yum123)* and *ilc-17*.*1 OE; nfki-1(yum127)*. Both *yum123* and *yum127* are null alleles of *nfki-1* generated by CRISPR. *ilc-17*.*1* extrachromosomal array was integrated into a locus tightly linked to *nfki-1* gene. Thus, two different alleles were used. Numerical values and statistical analyses are in Supplementary File 1. (**K**) Lifespan survival curves of WT and *ilc-17*.*1 OE* animals in 7% O_2_. Numerical values and statistical analyses are in Supplementary File 1.

**Figure 2–figure supplement 1.**
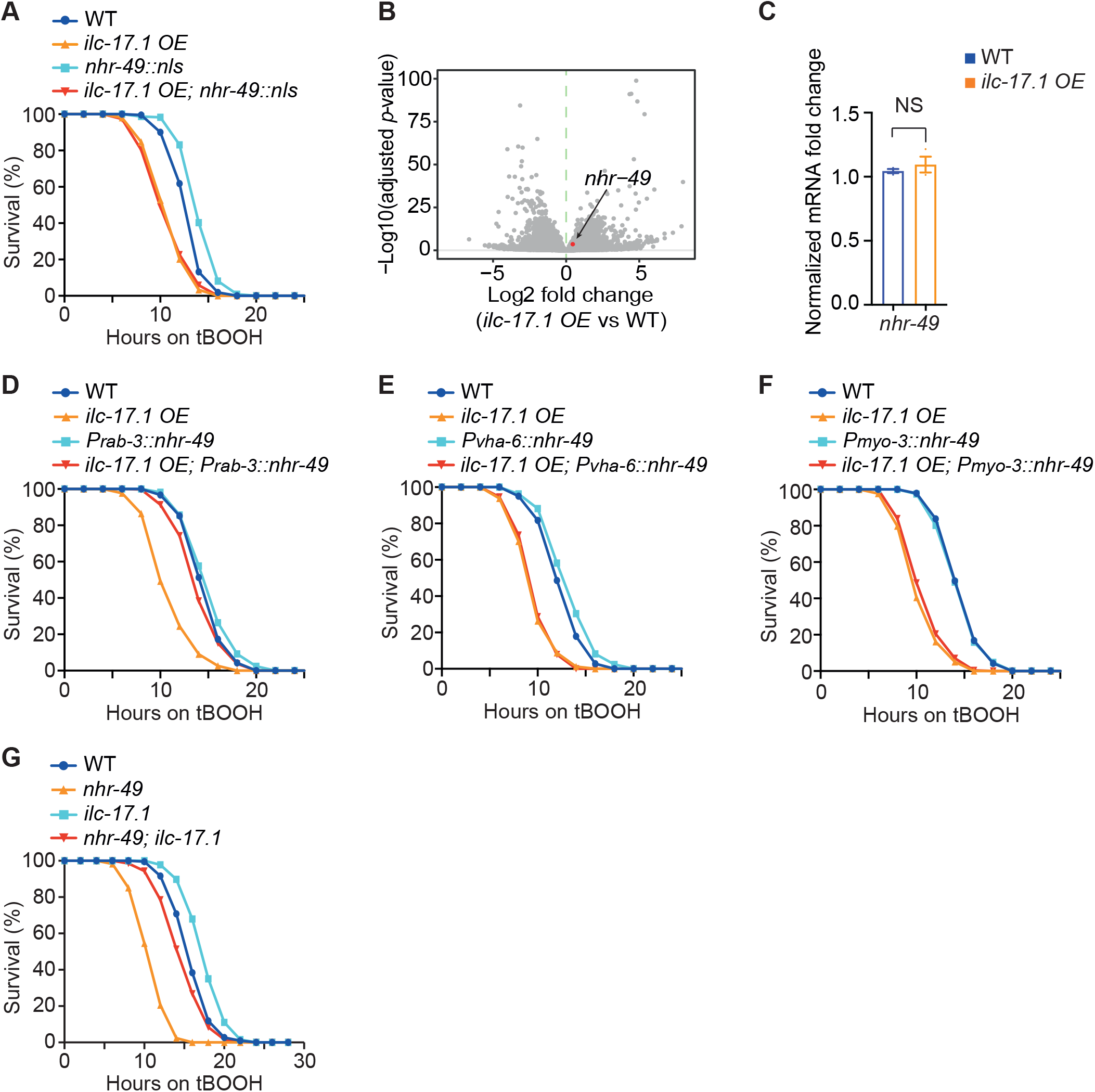
*nhr-49* transcript level is not altered by increased ILC-17.1 signaling. (**A**) Survival curves of WT, *ilc-17*.*1 OE, nhr-49::nls* and *ilc-17*.*1 OE; nhr-49::nls* animals on 7.7mM tBOOH. A SV40 nuclear localization signal (nls) was inserted before stop codon of *nhr-49* gene on the chromosome. Numerical values and statistical analyses are in Supplementary File 1. (**B**) Volcano plot highlighting the expression of *nhr-49* in *ilc-17*.*1 OE* animals relative to wild type. (**C**) Relative mRNA levels of *nhr-49* in WT and *ilc-17*.*1 OE* animals measured by qPCR. *act-1* and *tba-1* were used as the internal controls. Three independent experiments were performed, and each experiment contains 3 technical replicates. Error bars indicate SEM. NS = not significant, *t* test. (**D**) Survival curves of WT, *ilc-17*.*1 OE*, WT with *nhr-49* overexpressed in the neurons (*Prab-3*), and *ilc-17*.*1 OE* with *nhr-49* overexpressed in the neurons (*Prab-3*). Numerical values and statistical analyses are in Supplementary File 1. (**E**) Survival curves of WT, *ilc-17*.*1 OE*, WT with *nhr-49* overexpressed in the intestine (*Pvha-6*), and *ilc-17*.*1 OE* with *nhr-49* overexpressed in the intestine (*Pvha-6*). Numerical values and statistical analyses are in Supplementary File 1. (**F**) Survival curves of WT, *ilc-17*.*1 OE*, WT with *nhr-49* overexpressed in the muscle (*Pmyo-3*), and *ilc-17*.*1 OE* with *nhr-49* overexpressed in the muscle (*Pmyo-3*). Numerical values and statistical analyses are in Supplementary File 1. (**G**) Survival curves of WT, *nhr-49(nr2041), ilc-17*.*1(tm5218)* and *nhr-49(nr2041); ilc-17*.*1(tm5218)* animals on 7.7mM tBOOH. Numerical values and statistical analyses are in Supplementary File 1.

**Figure 3–figure supplement 1.**
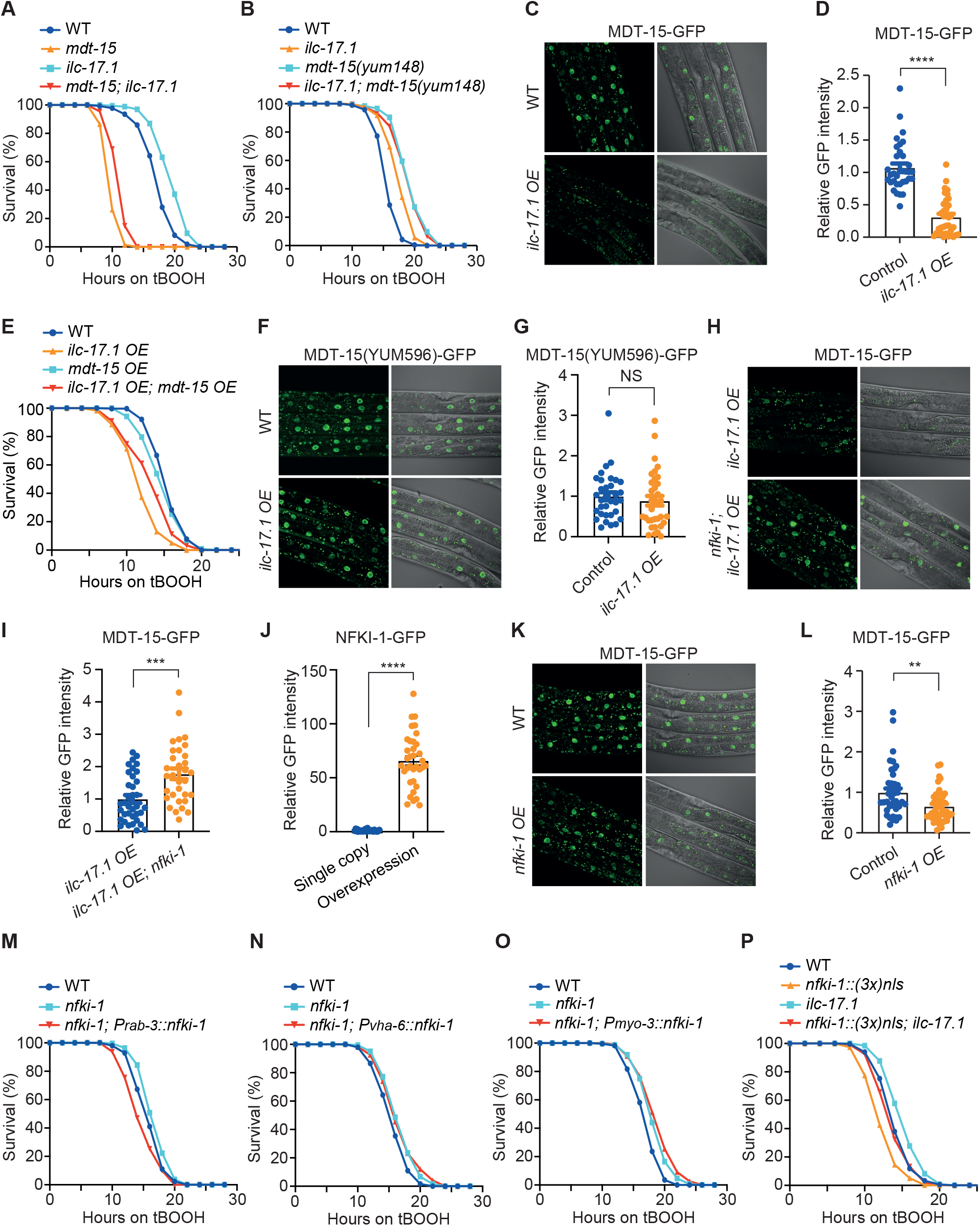
ILC-17.1 and MDT-15 act in the same pathway. (**A**) Survival curves of WT, *mdt-15(tm2182), ilc-17*.*1(tm5218)* and *mdt-15(tm2182); ilc-17*.*1(tm5218)* animals on 7.7mM tBOOH. Numerical values and statistical analyses are in Supplementary File 1. (**B**) Survival curves of WT, *ilc-17*.*1(tm5218), mdt-15(yum148)* and *ilc-17*.*1(tm5218)*; *mdt-15(yum148)* animals on 7.7mM tBOOH. *yum148* is a gain-of-function allele of *mdt-15*, equivalent to *mdt-15(et14)*. Numerical values and statistical analyses are in Supplementary File 1. (**C**) Representative images showing endogenously tagged MDT-15-GFP expression in the intestine of WT and *ilc-17*.*1 OE* animals. (**D**) Quantification of MDT-15-GFP fluorescent intensity in the intestinal nuclei of WT (n=34) and *ilc-17*.*1 OE* animals (n=35). Error bars indicate SEM. ***, *p*<0.001, *t* test. (**E**) Survival curves of WT, *ilc-17*.*1 OE, mdt-15 OE* and *ilc-17*.*1 OE*; *mdt-15 OE* animals on 7.7mM tBOOH. Numerical values and statistical analyses are in Supplementary File 1. (**F**) Representative images displaying endogenously tagged MDT-15(YUM596)-GFP expression in the intestine of WT and *ilc-17*.*1 OE* animals. *mdt-15(yum596)* is a gain-of-function allele of *mdt-15* generated by CRISPR, and is equivalent to *mdt-15(et14)*. (**G**) Quantification of MDT-15(YUM596)-GFP fluorescent intensity in the intestinal nuclei of WT (n=33) and *ilc-17*.*1 OE* animals (n=35). Error bars indicate SEM. NS = not significant, *t* test. (**H**) Representative images displaying endogenously tagged MDT-15-GFP expression in the intestine of *ilc-17*.*1 OE* and *ilc-17*.*1 OE; nfki-1(yum5066)* animals. *nfki-1(yum5066)* is a null allele generated by CRISPR. (**I**) Quantification of MDT-15-GFP fluorescent intensity in the intestinal nuclei of *ilc-17*.*1 OE* (n=36) and *ilc-17*.*1 OE; nfki-1(yum5066)* animals (n=32). Error bars indicate SEM. ***, *p*<0.001, *t* test. (**J**) Quantification of GFP intensity in the *nfki-1::gfp* single copy insertion (n=36) and *nfki-1::gfp OE* strains (n=35). *nfki-1::gfp* single copy was generated using CRISPR/Cas9, and *nfki-1::gfp OE* was generated by integrating *nfki-1::gfp* extrachromosomal arrays into the genome. The average fluorescent intensity of *nfki-1::gfp* single copy was arbitrarily set to 1. Error bars indicate SEM. ****, *p*<0.0001, *t* test. (**K**) Representative images showing endogenously tagged MDT-15-GFP expression in the intestine of WT and *nfki-1 OE* animals. (**L**) Quantification of MDT-15-GFP fluorescent intensity in the intestinal nuclei of WT (n=40) and *nfki-1 OE* animals (n=45). Error bars indicate SEM. **, *p*<0.01, *t* test. (**M**) Survival curves of WT, *nfki-1(yum123)* and *nfki-1(yum123)* with *nfki-1* overexpressed in the neurons (*Prab-3*). Numerical values and statistical analyses are in Supplementary File 1. (**N**) Survival curves of WT, *nfki-1(yum123)* and *nfki-1(yum123)* with *nfki-1* overexpressed in the intestine (*Pvha-6*). Numerical values and statistical analyses are in Supplementary File 1. (**O**) Survival curves of WT, *nfki-1(yum123)* and *nfki-1(yum123)* with *nfki-1* overexpressed in the muscle (*Pmyo-3*). Numerical values and statistical analyses are in Supplementary File 1. (**P**) Survival curves of WT, *nfki-1::(3x)nls, ilc-17*.*1(tm5218)*, and *nfki-1::(3x)nls; ilc-17*.*1(tm5218)* animals on 7.7mM tBOOH. Three copies of SV40 nuclear localization signals (3x *nls*) were inserted before stop codon of *nhr-49* gene on the chromosome. Numerical values and statistical analyses are in Supplementary File 1.

**Figure 4–figure supplement 1.**
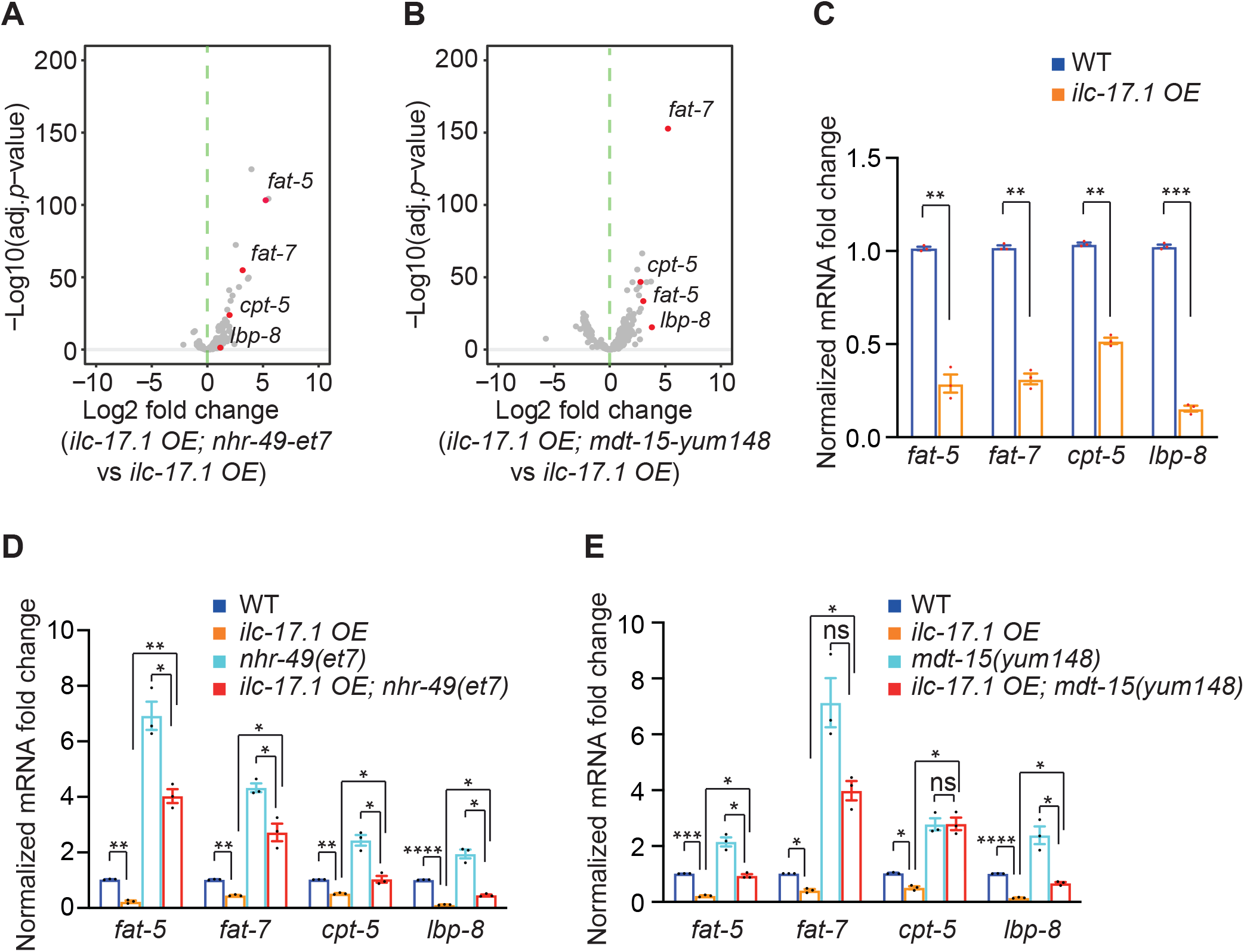
Relative mRNA expression of NHR-49/MDT-15 target genes. (**A**) Genes down-regulated by *ilc-17*.*1 OE* were included in the analysis. Volcano plot displaying the expression of these genes in *ilc-17*.*1 OE; nhr-49(et7)* relative to *ilc-17*.*1 OE* animals. The relative expression of *fat-5, fat-7, cpt-5* and *lbp-8* was highlighted. Please note that *cpt-5*, having adjusted *p*<1e-6, was not among 401 genes down-regulated by *ilc-17*.*1 OE* (adjusted *p*<1e-10), and was arbitrarily included in the analysis. (**B**) Genes down-regulated by *ilc-17*.*1 OE* were included in the analysis. Volcano plot displaying the expression of these genes in *ilc-17*.*1 OE; mdt-15(yum148)* relative to *ilc-17*.*1 OE* animals. The relative expression of *fat-5, fat-7, cpt-5* and *lbp-8* was highlighted. Please note that *cpt-5*, having adjusted *p*<1e-6, was not among 401 genes down-regulated by *ilc-17*.*1 OE* (adjusted *p*<1e-10), and was arbitrarily included in the analysis. (**C**) Relative mRNA levels of *fat-5, fat-7, cpt-5* and *lbp-8* in WT and *ilc-17*.*1 OE* animals measured by qPCR. *act-1* and *tba-1* were used as the internal controls. Three independent experiments were performed, and each experiment contains 3 technical replicates. Error bars indicate SEM. *, *p*<0.05, **, *p*<0.01, ***, *p*<0.001, *t* test. (**D**) Relative mRNA levels of *fat-5, fat-7, cpt-5* and *lbp-8* in WT, *ilc-17*.*1 OE, nhr-49(et7)* and *ilc-17*.*1 OE; nhr-49(et7)* animals measured by qPCR. *act-1* and *tba-1* were used as the internal controls. Three independent experiments were performed, and each experiment contains 3 technical replicates. Error bars indicate SEM. *, *p*<0.05, **, *p*<0.01, ***, *p*<0.001, *t* test. (**E**) Relative mRNA levels of *fat-5, fat-7, cpt-5* and *lbp-8* in WT, *ilc-17*.*1 OE, mdt-15(yum148)* and *ilc-17*.*1 OE; mdt-15(yum148)* animals measured by qPCR. *act-1* and *tba-1* were used as the internal controls. Three independent experiments were performed, and each experiment contains 3 technical replicates. Error bars indicate SEM. NS = not significant, *, *p*<0.05, **, *p*<0.01, ***, *p*<0.001, *t* test.

**Figure 5–figure supplement 1.**
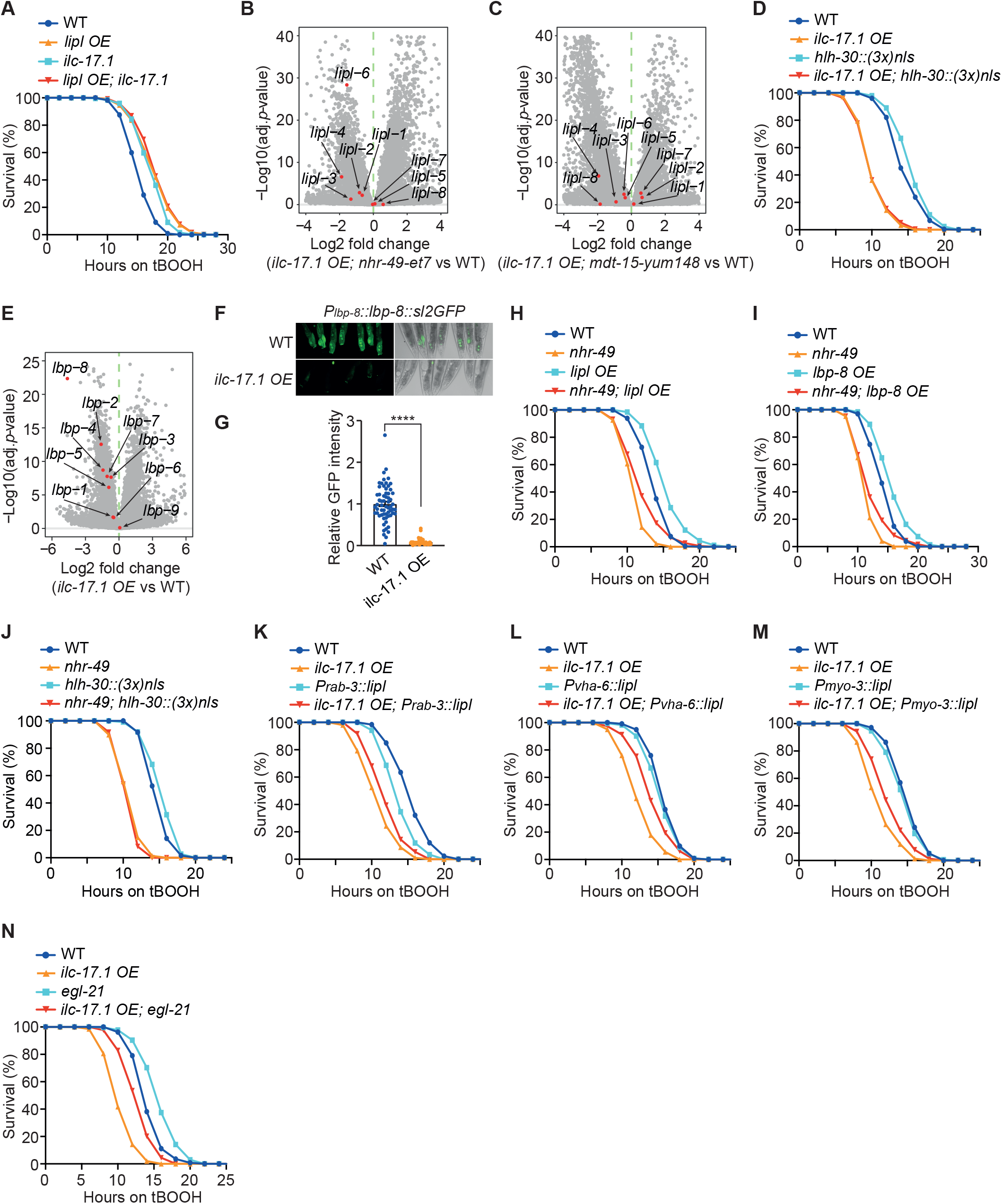
Relative mRNA expression of *lipl* and *lbp* genes. (**A**) Survival curves of WT, *lipl OE, ilc-17*.*1(tm5218)*, and *lipl OE; ilc-17*.*1(tm5218)* animals on 7.7mM tBOOH. *lipl OE* is the overexpression of *lipl-1, lipl-2, lipl-3* and *lipl-4* from extrachromosomal arrays. Numerical values and statistical analyses are in Supplementary File 1. (**B** and **C**) Volcano plots highlighting the expression of *lipl* genes in *ilc-17*.*1 OE; nhr-49(et7)* (B) and in *ilc-17*.*1 OE; mdt-15(yum148)* (C) animals relative to WT. (**D**) Survival curves of WT, *ilc-17*.*1 OE, hlh-30::(3x)nls*, and *ilc-17*.*1 OE*; *hlh-30::(3x)nls* animals on 7.7mM tBOOH. Three copies of SV40 nuclear localization signals (3x *nls*) were inserted before stop codon of *hlh-30* gene on the chromosome. Numerical values and statistical analyses are in Supplementary File 1. (**E**) Volcano plot highlighting the expression of *lbp* genes in *ilc-17*.*1 OE* animals relative to WT. (**F**) Representative images showing *gfp* expression driven by the promoter of *lbp-8* in WT and *ilc-17*.*1 OE* animals. (**G**) Quantification of GFP expression driven by *lbp-8* promoter in WT (n=62) and *ilc-17*.*1 OE* animals (n=49). ****, *p*<0.0001, *t* test. (**H**) Survival curves of WT, *nhr-49(nr2041), lipl OE* and *nhr-49(nr2041); lipl OE* animals on 7.7mM tBOOH. *lipl OE* is the overexpression of *lipl-1, lipl-2, lipl-3* and *lipl-4* from extrachromosomal arrays. Numerical values and statistical analyses are in Supplementary File 1. (**I**) Survival curves of WT, *nhr-49(nr2041), lbp-8 OE* and *nhr-49(nr2041); lbp-8 OE* animals on 7.7mM tBOOH. Numerical values and statistical analyses are in Supplementary File 1. (**J**) Survival curves of WT, *nhr-49(nr2041), hlh-30::(3x)nls*, and *nhr-49(nr2041)*; *hlh-30::(3x)nls* animals on 7.7mM tBOOH. Three copies of SV40 nuclear localization signals (3x *nls*) were inserted before stop codon of *hlh-30* gene on the chromosome. Numerical values and statistical analyses are in Supplementary File 1. (**K**) Survival curves of WT, *ilc-17*.*1 OE, lipl* genes overexpressed in the neurons (*Prab-3*), and *ilc-17*.*1 OE* with *lipl* genes overexpressed in the neurons (*Prab-3*). Numerical values and statistical analyses are in Supplementary File 1. (**L**) Survival curves of WT, *ilc-17*.*1 OE, lipl* genes overexpressed in the intestine (*Pvha-6*), and *ilc-17*.*1 OE* with *lipl* genes overexpressed in the intestine (*Pvha-6*). Numerical values and statistical analyses are in Supplementary File 1. (**M**) Survival curves of WT, *ilc-17*.*1 OE, lipl* genes overexpressed in the muscle (*Pmyo-3*), and *ilc-17*.*1 OE* with *lipl* genes overexpressed in the muscle (*Pmyo-3*). Numerical values and statistical analyses are in Supplementary File 1. (**N**) Survival curves of WT, *ilc-17*.*1 OE, egl-21(n476)* and *ilc-17*.*1 OE; egl-21(n476)* animals on 7.7mM tBOOH. Numerical values and statistical analyses are in Supplementary File 1.

**Figure 6–figure supplement 1.**
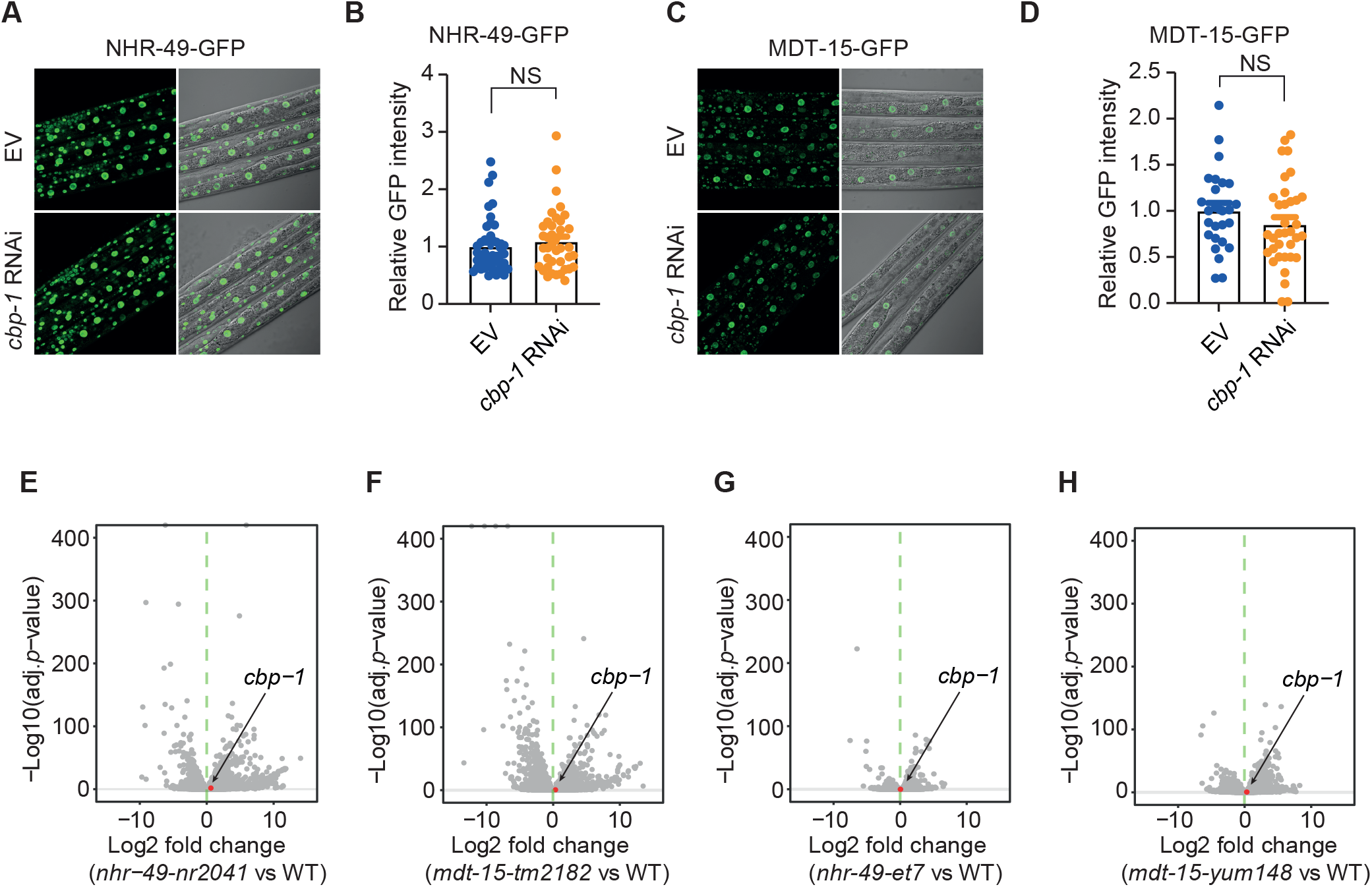
The expression of NHR-49/MDT-15 and CBP-1. (**A**) Representative images displaying endogenously tagged NHR-49-GFP expression in the WT animals treated with either RNAi empty vector (EV) L4440 (control) or a RNAi plasmid targeting *cbp-1. cbp-1* RNAi culture was diluted (1:20) by the control before seeding on the RNAi plates. (**B**) Quantification of NHR-49-GFP intensity in the WT animals treated with either RNAi empty vector (EV) L4440 (control) (n=44) or a RNAi plasmid targeting *cbp-1. cbp-1* RNAi culture was diluted (1:20) by the control before seeding on the RNAi plates (n=39). Error bars indicate SEM. NS = not significant, *t* test. (**C**) Representative images displaying endogenously tagged MDT-15-GFP expression in the WT animals treated with either RNAi empty vector (EV) L4440 (control) or a RNAi plasmid targeting *cbp-1. cbp-1* RNAi culture was diluted (1:20) by the control before seeding on the RNAi plates. (**D**) Quantification of MDT-15-GFP intensity in the WT animals treated with either RNAi empty vector (EV) L4440 (control) (n=26) or a RNAi plasmid targeting *cbp-1. cbp-1* RNAi culture was diluted (1:20) by the control before seeding on the RNAi plates (n=35). Error bars indicate SEM. NS = not significant, *t* test. (**E**) Volcano plot highlighting the expression of *cbp-1* in *nhr-49(nr2041)* animals relative to wild type. (**F**) Volcano plot highlighting the expression of *cbp-1* in *mdt-15(tm2182)* animals relative to wild type. (**G**) Volcano plot highlighting the expression of *cbp-1* in *nhr-49(et7)* animals relative to wild type. (**H**) Volcano plot highlighting the expression of *cbp-1* in *mdt-15(yum148)* animals relative to wild type.

